# Molecular Systems Predict Equilibrium Distributions of Phenotype Diversity Available for Selection

**DOI:** 10.1101/2021.05.27.446045

**Authors:** Miguel A. Valderrama-Gómez, Michael A. Savageau

## Abstract

Two long standing challenges in theoretical population genetics and evolution are predicting the distribution of phenotype diversity generated by mutation and available for selection and determining the interaction of mutation, selection, and drift to characterize evolutionary equilibria and dynamics. More fundamental for enabling such predictions is the current inability to causally link population genetic parameters, selection and mutation, to the underlying molecular parameters, kinetic and thermodynamic. Such predictions would also have implications for understanding cryptic genetic variation and the role of phenotypic robustness.

Here we provide a new theoretical framework for addressing these challenges. It is built on Systems Design Space methods that relate system phenotypes to genetically-determined parameters and environmentally-determined variables. These methods, based on the foundation of biochemical kinetics and the deconstruction of complex systems into rigorously defined biochemical phenotypes, provide several innovations that automate (1) enumeration of the phenotypic repertoire without knowledge of kinetic parameter values, (2) representation of phenotypic regions and their relationships in a System Design Space, and (3) prediction of values for kinetic parameters, concentrations, fluxes and global tolerances for each phenotype.

We now show that these methods also automate prediction of phenotype-specific mutation rate constants and equilibrium distributions of phenotype diversity in populations undergoing steady-state exponential growth. We introduce this theoretical framework in the context of a case study involving a small molecular system, a primordial circadian clock, compare and contrast this framework with other approaches in theoretical population genetics, and discuss experimental challenges for testing predictions.

## INTRODUCTION

The concept of evolution is easily stated and understood: Mutation generates diversity of phenotypes and selection favors those with the greatest heritable fitness. However, there are many complex and inter-related issues that must be addressed to achieve a deeper understanding. Two prominent examples that continue to be fundamental challenges are (1) determining the *distribution of phenotype diversity*, which offers opportunities for innovation (Charlesworth, 1996; Bataillon & Bailey, 2014) and (2) determining the interaction of *mutation, selection, drift and population structure* to determine equilibria and the dynamics of evolution (Gillespie, 2004; Orr, 2005; Wakeley, 2005).

While the distribution in the numbers and types of changes in DNA can be determined because of advances in genome sequencing technology (Metzker, 2010), determining the distribution of the resulting phenotypes and their fitness characteristics (determinants of total fitness) in natural populations is difficult in the extreme (Charlesworth, 1996). The true number of phenotypes and fitness characteristics in the population is typically unknown and any observed distribution of total fitness (e.g., growth rate of bacteria) is skewed by what can be observed in the field or measured or generated experimentally in the laboratory (Gallet et al., 2012; Robert et al., 2018; Bondel et al., 2019; Lebeuf-Taylor, et al., 2019). With large data sets, correlations can be established between genome changes and fitness changes in a given environment. However, at a fundamental level there are many fitness components based on function that remain to be identified and characterized and many unsolved mappings that prevent a predictive, causal linking of mutations in DNA, properties of molecular components, integrated system function, phenotypic repertoire, and fitness. In short, there is little relevant theory for guidance.

There is a rich field of theoretical population genetics developed over more than a century that addresses the interaction of *mutation, selection, drift and population structure* (Gillespie, 2004; Orr, 2005; Wakeley, 2005). However, aside from the simpler cases of one-to-one mapping between gene and phenotypic function, an appropriate theoretical framework is lacking to pose and answer questions for the more complex cases that involve mappings between many genes and many functional contributions to phenotypes.

In reviewing the genetic theory of adaptation, Orr (2005) examined “the reasons a mature [mathematical] theory has been slow to develop and the prospects and problems facing current theory” and concluded that although recent models “seem to successfully explain certain qualitative patterns […] future work must determine whether present theory can explain the genetic data quantitatively”. Experimental evolution studies have shown that mutations in a single gene affecting a specific enzyme can lead to a marked change in organismal fitness (Barrick & Lenski, 2013; Gresham & Jong, 2015). Although the results might be explained qualitatively, without an adequate systems theory these explanations cannot provide a rigorous, quantitative, causal understanding of the complex underlying events.

Can knowledge of molecular systems tell us anything about the distribution of mutant phenotypes and their evolution? A large part of the problem in relating molecular mechanisms to phenotype distributions and evolution is the inability to relate the genotype and environment to the phenotype exhibited by a biological system, which is one of the ‘Grand Challenges’ in biology (Brenner, 2000). The causal linking of genotype to phenotype involves at least three essential mappings: First is the mapping from the *digital values* of the genome sequence to the *analogue values* of the kinetic parameters that characterize the underlying molecular processes. Second is the mapping from the *kinetic parameters* of the individual component processes to the quantitative *biochemical phenotypes* of the integrated cellular system. Third is the mapping from the *biochemical* (endo-) *phenotypes* to the *organismal (exo-) phenotypes*, including observables such as growth rate, taxis, adhesion, etc. The first of these mappings deals with protein structure function relationships, which relate DNA sequence to properties of the encoded protein. Recent success in solving the protein folding problem (Callaway, 2020) bodes well for the eventual ability to predict kinetic parameters. The second mapping is the focus of Biochemical Systems Theory (Savageau, 1971; 2009; Voit, 2000; 2013), which in the past decade has provided a novel system deconstruction that maps genetically-determined parameters and environmentally-determined variables to biochemical phenotypes. The result is a highly structured partitioning of parameter space that is defined as the *System Design Space* (Savageau et al., 2009). A Design Space Toolbox (DST3) is available with numerous tools that automate the analysis (Valderrama-Gómez et al., 2020). The third of these mappings is perhaps the most difficult, in any but the simplest cases of one-gene one-protein one-phenotype, due to the large number of genes and phenotypes with many-to-many interactions that currently can only be characterized by large data sets and statistical correlations (McCarthy et al., 2008; Greenbury et al., 2016).

If we could enumerate the full repertoire of phenotypic functions that could be exhibited by a given biological system and know the ranking of phenotype frequencies in the population undergoing mutational exchange, we would have a deep understanding of the functional basis for phenotypic diversity and plasticity available for selection to act upon. Here we address these issues in five parts.

First, we introduce a small molecular system, a primordial precursor to a circadian clock, as an aid to understanding this novel approach. We have specifically selected a hypothesis-motivated example with unknown parameter values for this purpose because any real system will initially have many unknowns and involve the formulation of, and discrimination among, many hypotheses that require experimental testing. Our approach has advantages to offer specifically at this stage of an investigation (Lomnitz & Savageau, 2016a). Second, we utilize System Design Space concepts **(Supplemental Information, Section S1)** to extend and apply the phenotype-centric strategy to predict phenotype-specific mutation rate constants. This involves formulating phenotype-specific mutation rates based on transition probabilities between biochemical phenotypes. These rate constants are then used to formulate population dynamic equations for predicting equilibrium distributions of phenotype diversity under non-selecting and selecting conditions. Third, results for the population genetic model are presented. Fourth, specific predictions are discussed in the light of experimental challenges for their testing. Fifth, in the discussion, we briefly compare and contrast the theoretical framework provided by the System Design Space approach with that provided by other approaches.

## PUTATIVE PRIMORDIAL CIRCADIAN CLOCK

The molecular system, treated as a case study here, is related to the positive-negative feedback module found at the core of nearly all circadian clocks (Bell-Pedersen et al., 2005; Hardin, 2011; Cohen & Golden, 2015; Nohales & Kay, 2016; Papazyan et al., 2016; Creux & Harmer, 2020) and several synthetic oscillator designs (Atkinson et al., 2003; Stricker et al., 2008; Tigges et al., 2009; Lomnitz & Savageau, 2014). In the transcription-translation oscillators, this module consists of a positive transcription factor that activates its own synthesis as well as synthesis of a negative transcription factor, which in turn represses synthesis of the positive transcription factor. The originally identified module in *Drosophila* is elaborated upon in animals (Preitner etal., 2002) and plants (Creux & Harmer, 2020) with numerous variations on the theme, including diverse input stimuli that modulate expression of one or both factors (Balsalobre, et al. 2000; O’Neill & Reddy, 2012) and rich output interactions with nearly all cellular functions (Creux & Harmer, 2020).

In the cyanobacterial clock, the transcription-translation mechanism is a minor player whereas a posttranslational oscillator mechanism with different positive and negative interactions plays the dominant role (Cohen & Golden, 2015). When growing exponentially in a normal diurnal light cycle, phenotypes without the oscillatory characteristic are at a selective disadvantage when compared to the wild type (oscillatory phenotype); however, they exhibit no measurable disadvantage when grown under the non-selecting condition (constant light), as determined by growth competition between mutants and wild type in an otherwise isogenic background (Ouyang et al., 1998). Under these conditions, biochemical phenotypes map closely to organismal phenotypes, which is the assumption typically made in studying the evolution of specific molecular systems in bacteria (Brajesh, et al., 2019).

Roenneberg & Merrow (2002) and many others have speculated that the robust limit cycle or sustained oscillation exhibited by circadian clocks in modern organisms is unlikely to have arisen full blown. We propose that some of the coordinating functions could have been provided by a simpler core module having a damped oscillation with a frequency that resonates to and becomes synchronized with the diurnal cycle. Indeed, such damped oscillations have been experimentally observed in strains of cyanobacteria: namely, clock mutants of *Synechococcus* (Ouyang et al., 1998; Kawamoto et al., 2020) and marine *Prochlorococcus marinus* (Holtzendorff et al., 2008).

For our purposes here, we shall consider the primordial mechanisms to involve only the negative feedback loop at the core of modern transcription-translation mechanisms, as shown schematically in **Figure 1**. The equations used to represent this putative primordial clock (**Supplemental Information, Section S2**) are based on the foundation of fundamental biochemical kinetics. These models have broad general applicability, as the vast majority of biochemical models are of this type (Chelliah et al., 2013). These equations, recast as an equivalent GMA-system of equations (**Supplemental Information, Section S3**), can be expressed as a set of differential-algebraic equations in the syntax of the Design Space Toolbox (Lomnitz & Savageau, 2016b).

**Figure 1.**
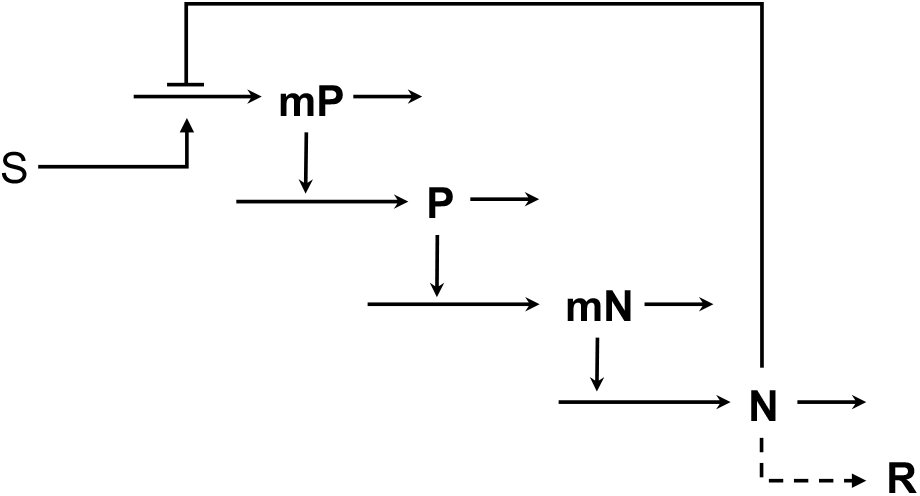
Common genetic module for the putative precursor of the modern core mechanism of nearly all circadian clocksxsx. Positive (P) and negative (N) transcription factor proteins and the corresponding mRNAs (mP and mN). Environmental input stimulus (S) and biochemical output response (R) are suggestive only, other targets and coordinating signals could be considered. See also **Supplemental Information, Section S2**.

For the illustrative purposes of this case study, we make three simplifying assumptions. (1) The precursor is likely to involve a minimal number of processes and a minimal degree of cooperativity in the interactions. The two-transcription factor model involves at least four processes and two cooperative DNA interactions. For it to generate a damped oscillatory response, the system must be near the threshold of instability, which requires a value of loop cooperativity [(*n*p*) in the **Eqn**. (**S1**) to **Eqn**. (**S4**)] equal to 4 for a system with four temporally dominant stages (Savageau, 1975; Thron, 1991). We let the cooperativity parameters *n=p*=2. Kawamoto et al. (2020) considered a simpler three-stage model for *Synechococcus*; but it requires a much higher degree of cooperativity, *n* >8 as shown in Savageau (1975). (2) To provide the most challenging shape for testing different methods of volume calculation, we select values for the two parameters (capacity for regulation for the two transcripts) with the potential to break the symmetry such that a skewed volume is generated for the phenotype with an oscillatory characteristic (**Supplemental Information, Section S4, Figure S1**). (3) To aid visualization of the results we focus on a two-dimensional slice through the System Design Space with the two binding constants displayed on the vertical and horizontal axes, which is representative of the invariant for this system design space (**Supplemental Information, Section S5, Figure S2**).

The Design Space Toolbox 3 (DST3, Valderrama-Gómez et al., 2020) can be used to enumerate the repertoire of phenotypes without assuming values for any of the model’s kinetic parameters, and the results demonstrate a maximum of nine possible phenotypes. The listing of the phenotypes, along with the properties of their eigenvalues when *n=p*=2, is shown in **Table 1**. Each sequential pair of integers in the phenotype signature identifies the specific positive and negative terms in the corresponding GMA equation that are instrumental in defining the phenotype. A comparison of the phenotype signatures with the GMA equations in **Supplemental Information, Section S3**, identifies the specific S-system equation for each phenotype. All phenotypes are stable with no complex conjugate eigenvalues, except for phenotype #7; thus, only phenotype #7 has the potential to initiate damped oscillatory behavior.

**Table 1.**
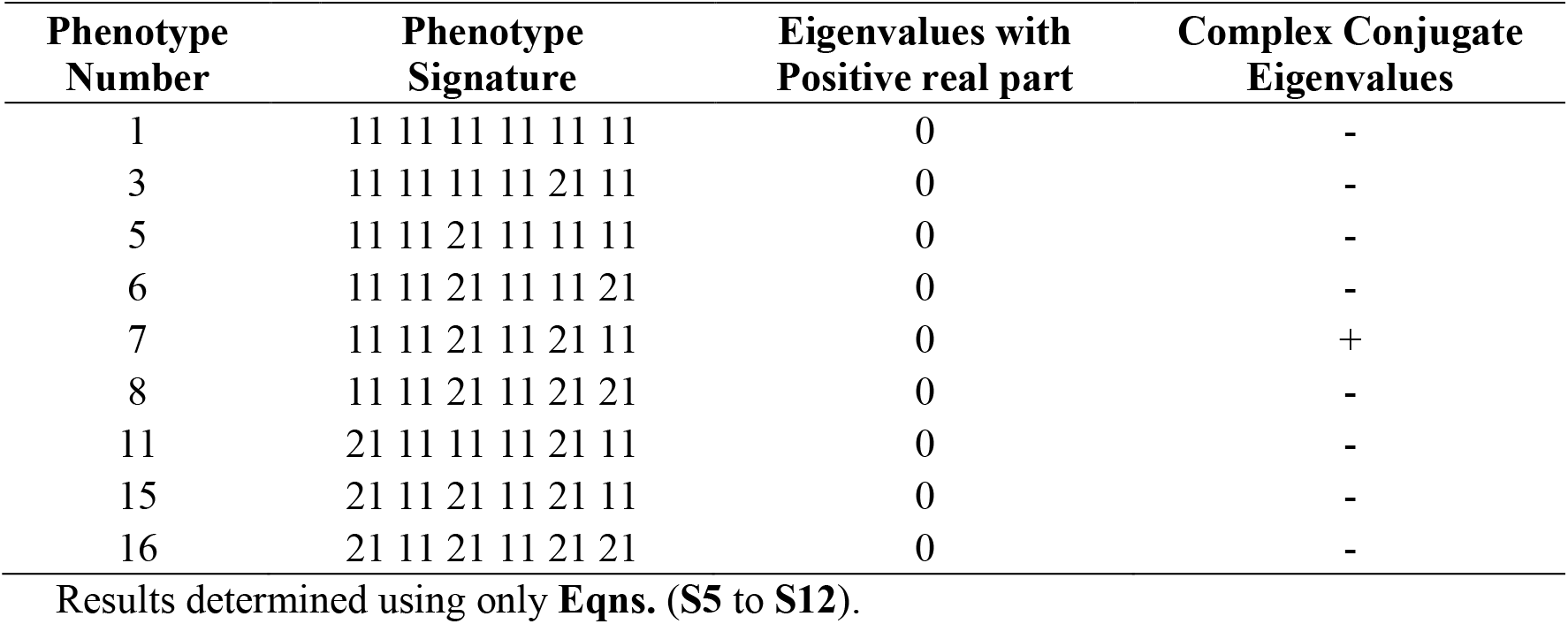
Phenotypic repertoire for the model in Figure 1.

| Phenotype Number | Phenotype Signature | Eigenvalues with Positive real part | Complex Conjugate Eigenvalues |
| --- | --- | --- | --- |
| 1 | 11 11 11 11 11 11 | 0 | - |
| 3 | 11 11 11 11 21 11 | 0 | - |
| 5 | 11 11 21 11 11 11 | 0 | - |
| 6 | 11 11 21 11 11 21 | 0 | - |
| 7 | 11 11 21 11 21 11 | 0 | + |
| 8 | 11 11 21 11 21 21 | 0 | - |
| 11 | 21 11 11 11 21 11 | 0 | - |
| 15 | 21 11 21 11 21 11 | 0 | - |
| 16 | 21 11 21 11 21 21 | 0 | - |

Results determined using only **Eqns. (S5 to S12)**.

It should be emphasized that the enumeration of the full repertoire by DST3 is accomplished *without having to specify values for any of the thermodynamic and kinetic parameters*. By specifying the stoichiometry for binding repressor and activator as *n*=2 and *p*=2, DST3 automatically predicts scaled values (**Table 2**, Legend) of all 12 thermodynamic and kinetic parameter values of the system, identifies the region in design space for the realization of phenotype #7, the phenotype of interest here, as well as the steady-state values of the four dynamic variables. By choosing the simplest scaling, generating a skewed volume for phenotype #7, and shifting the entire Design Space to center the visualization on phenotype #7 (**Supplemental Information, Section S5, Figure S2**), we predict values for the 12 parameters and the steady-state values for the four dynamic variables as shown in **Table 2**.

**Table 2.**
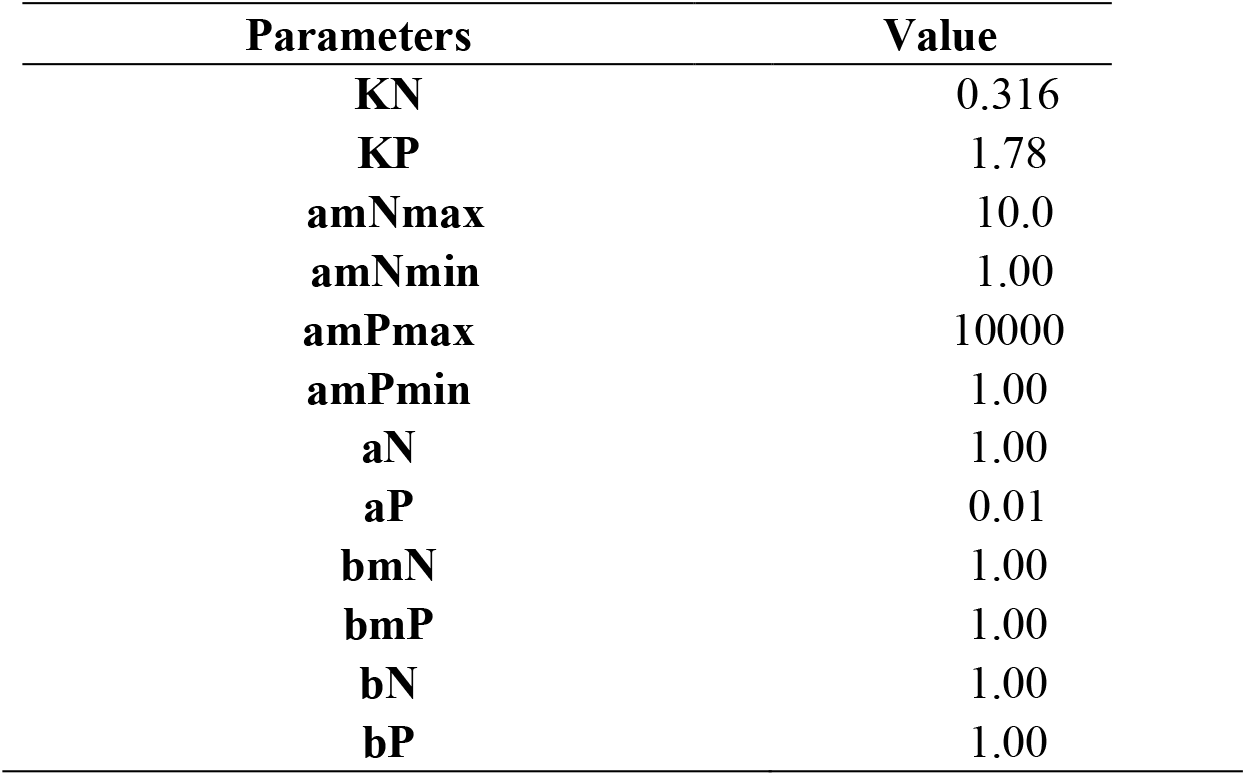
Scaled values for the parameters and steady state concentrations automatically determined and fixed for phenotype 7 (11 11 21 11 21 11). The behavior of the model is determined by these scaled parameter values. If necessary, twelve experimental measurements (e.g., maximum expression, minimum expression and lifetime of each mRNA and protein) are sufficient to determine the actual parameter values. However, as can be seen in the **Eqn. (1)**, our methods involve differences in log space so the scale factors cancel out and thus there is no effect on the qualitative or quantitative results. Predicted normalized steady-state values: mP = 100.0; P = 1.0; mN = 3.16; N = 3.16.

| Parameters | Value |
| --- | --- |
| **KN** | 0.316 |
| **KP** | 1.78 |
| **amNmax** | 10.0 |
| **amNmin** | 1.00 |
| **amPmax** | 10000 |
| **amPmin** | 1.00 |
| **aN** | 1.00 |
| **aP** | 0.01 |
| **bmN** | 1.00 |
| **bmP** | 1.00 |
| **bN** | 1.00 |
| **bP** | 1.00 |

Although we could examine variations in all 12 parameters, we choose to fix the predicted values for all parameters except for the two equilibrium dissociation constants *K*_*P*_ and *K*_*N*_, which will be allowed to vary because of mutation. This simplification reduces the dimensions of the Design Space for ease in visualizing the results while providing an accurate representation of the underlying Design Space invariant. The size of the regions in Design Space occupied by each of the phenotypes (**Figure 2A**) then can be determined by a vertex enumeration method (Avis 2000, Barber et al. 1996). These methods work well for small systems and other methods are available for large systems (**Supplemental Information, Section S6, Figure S3**).

**Figure 2.**
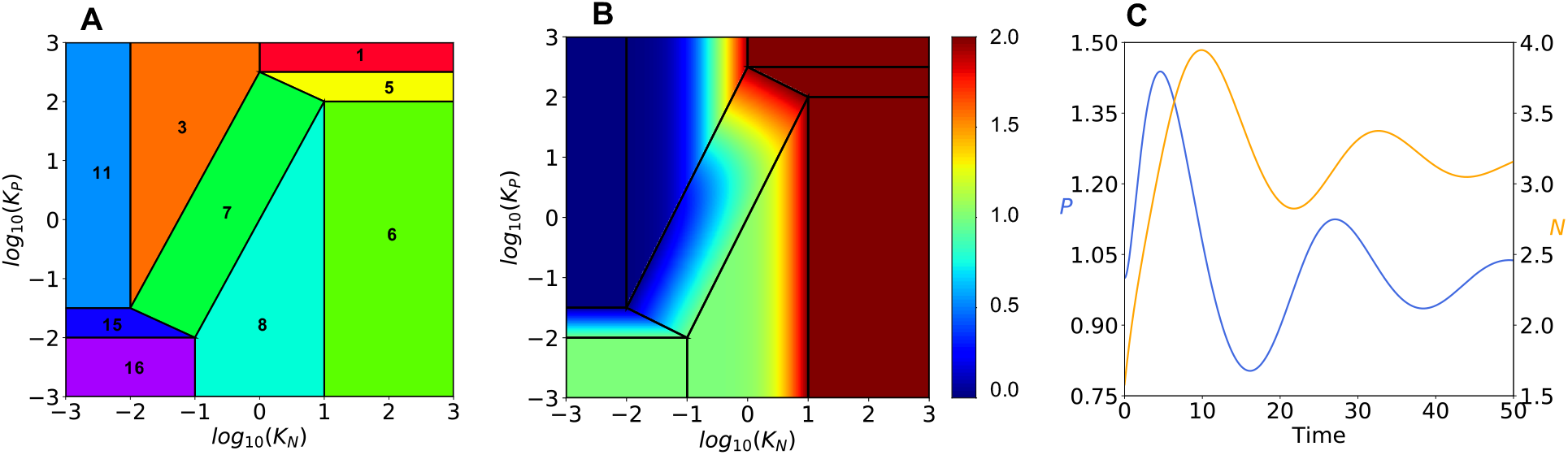
Predicted phenotype characteristics in System Design Space. **(A)** Visualization of phenotype regions. Region of oscillatory phenotype #7 is the central rectangular shape. **(B)** Steady state concentration of total protein (N+P) plotted log10 as a heat map on the z-axis. **(C)** Validated oscillatory behavior for phenotype #7. Concentrations of activator P (left y-axis, Blue) and repressor N (right y-axis, Gold) as a function of time scaled by a factor of 1/3. Initial conditions are: mP=100; P = 1.0; mN = 3.16; N = 1.58. Figures generated with the following parameter values: KN = 0.316; KP = 1.78; aN = 1.0; aP = 0.01; amNmax = 10.0; amNmin = 1.0; amPmax = 10000.0; amPmin = 1.0; bN = 1.0; bP = 1.0; bmN = 1.0; bmP = 1.0; Kinetic order(s): n=2, p=2; (The parametric constraints amPmax>amPmin and amNmax>amNmin are automatically satisfied by this parameterization of the model.)

We are fully aware that this simplified example used to introduce our theoretical framework does not illustrate its full potential. There is a growing number of molecular systems for which there is information sufficient to specify their architecture. By model architecture we mean molecules, interactions among them, and the signs of the interactions. Surprisingly, much of what can be learned about the system depends largely on its architecture. It suggests mechanistic hypotheses without having to know details of the many kinetic and thermodynamic parameters. As noted in the **INTRODUCTION**, this is all that is needed to formulate testable hypotheses amenable to analysis in the System Design Space framework. One the other hand, we are also fully aware that this simplified example does not address numerous realistic and important questions in the large field of theoretical population genetics. These issues must be the subject of future work.

## NEW APPROACHES: DERIVATION OF PHENOTYPE-SPECIFIC MUTATION RATE CONSTANTS AND POPULATION DYNAMIC EQUATIONS

The System Design Space approach enables a novel ‘phenotype-centric’ modeling strategy that is radically different from the conventional ‘simulation-centric’ approach (Valderrama-Gómez et al., 2018). A summary of System Design Space concepts is provided (**Supplemental Information, Section S1)** to facilitate understanding of the phenotype-centric strategy used to predict phenotype-specific mutation rate constants.

The mechanistic framework we are proposing requires new concepts and methods; the reader is directed to **Supplemental Information** where these are fully developed. Here we simply summarize the four factors in System Design Space contributing to the probability of transition between phenotypes by mutational change in the mechanistic parameters: (1) mutated parameters change value along ‘tracks’ orthogonal to non-mutated parameters, (2) ‘volume’ of the phenotype resulting from mutation, (3) ‘size scale’ of parameter changes between original (doner) and resultant (recipient) phenotypes, and (4) ‘directional bias’ of parameter changes that are more probable in one direction vs. the alternative. These four factors are elaborated on in the following sections and it will be most helpful to visualize these factors in terms of the geometry of System Design Space that is determined by the architecture of the underlying molecular system (**Supplemental Information, Section S7**).

### Mutated vs. Orthogonal Parameters

We consider only mutational events that influence a single parameter. Thus, we need only consider a single ‘track’ for the mutated parameter between values of orthogonal parameters. Such mutations can influence multiple systemic functions indirectly, which is pleiotropy at the phenotype level. The tracks are rigorously defined by the vertices of the phenotype polytopes and, in some cases, polytopes are split into more than one track (**Figure S4D**). The single parameter restriction can be relaxed to consider mutations that influence multiple parameters of a single component, which might be considered pleiotropy at the single molecule level (e.g., the degradation rate constant of a transcription factor and its DNA binding constant). The single parameter restriction can also be relaxed to consider simultaneously multiple mutants (e.g., rare double mutant events). However, to make causal predictions upon removal of the single parameter restriction will require formulation of testable mechanistic hypotheses (Lomnitz & Savageau, 2016a).

### Phenotype Volume

Because the volume of a phenotype becomes infinite when the phenotype is independent of some parameter in the model, we bound the universe of values for all parameters by a hyper-cube in log space that is Π-orders of magnitude on edge. The value of Π should be large enough to include all phenotypes in the System Design Space but not so large as to exceed physically realistic parameter values; this will exclude phenotypes that can only be realized with unrealistic parameter values. We have set Π = 6, which seems large enough to cover all values that can be distinguished experimentally, which is typically about 3-orders of magnitude. For example, the repressor for the lactose operon of *Escherichia coli* binds tightly to specific recognition sites in the DNA with an occupancy of nearly 100%, but reduction of its equilibrium dissociation constant by three orders of magnitude reduces the occupancy to nearly 0% (Lewin, 2008). Moreover, it is very unlikely that parameter values ever go to zero because there are typically promiscuous proteins capable of performing the same function with at least some minimal activity (Khersonsky & Tawfin, 2010; Rueda et al., 2019). In any case, we have obtained similar results with Π = 8, and DST3 allows users to select a custom value for Π.

Given a particular set of parameter values characterizing the donor phenotype in volume *V*_*i*_, one of the four contributions to the probability of mutating to any other set of parameter values in the volume of the recipient phenotype *V*_*j*_, along a given mutant parameter track, is given by the ratio of the recipient volume to the total volume along the entire track. Thus, this contribution to the probability of mutating from a phenotype with a small track volume to one with a large track volume is greater than in the opposite direction.

### Size Scale

Large scale mutations are rare; small scale mutations are frequent in well adapted systems (Bataillon & Bailey, 2014; Tataru et al., 2017; Bondel et al., 2019; Templeton, 2021). This size-scale effect depends on the distance, *s*, between the operating point (a parameter set) of the donor phenotype and that of the recipient phenotype. By sampling each donor and recipient combination along the line representing the change in the mutated mechanistic parameter, the probability of each mutation can be calculated based on the track volume of the recipient phenotype and distribution of size-scale effects for the mutations. Although the actual distributions for size-scale effect are unknown, we assume that the probability of parameter change by mutation decreases exponentially with a size scale λ, i.e. ∼ exp(-*s/λ*). This will be made more concrete in **RESULTS (*first subsection*)**.

The size-scale effect of mutations can be calculated as an average distance over all combinations of donor and recipient parameter values within a track, which is computationally demanding, or by considering the distance between ‘phenotype centroids’ (red dots in **Figure S4B**,**C**,**D**), analogous to the ‘centers of mass’ in mechanics. The results are the same for both methods (**Supplemental Information, Section S8, Figure S5**) and, since it is computationally more efficient, we use the centroid method.

### Directional Bias

The probability of mutation can be further refined by consideration of “directional bias”. The probability is larger when a parameter change is in the direction of increasing entropy; it is smaller when the change is in the direction of decreasing entropy. Although the actual differences in value are currently unknown, we account for these directional biases by assigning a multiplicative weighting factor δ that increases the effective size scale λ when a parameter change is in the direction of increasing entropy and decreases it when the parameter change is in the direction decreasing entropy (**Supplemental Information, Section S9, Table S1**).

### Mutation Rates

*Phenotype-specific mutation rate constants* are determined in three steps. First, for each donor *i* and recipient *j* phenotype, the mechanistic parameter contribution to the mutation, ***K***_*ij*_, is determined with an exponential distribution (**Table S1**) involving size scale *λ*, directional bias *δ*, and the magnitude of parameter difference *s*, between phenotype centroids *C*_*i*_, i.e.,

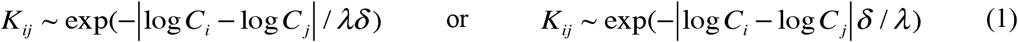

depending on whether the directional bias of the mutation is toward increased entropy (1 / ***δ***) or decreased entropy (***δ***). The product of the mechanistic contribution, ***K***_*ij*_, and the track volume of the recipient phenotype, *V*_*j*_, is proportional to the probability of a mutation from donor phenotype *i* to recipient phenotype *j*.

Second, the normalized probability of a mutation from donor phenotype *i* to recipient phenotype *j* along a track variable is written

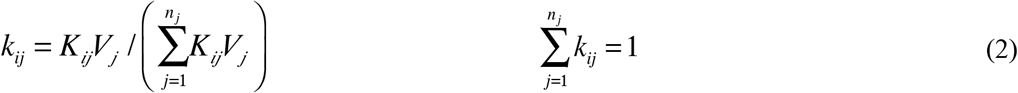

where *n*_*j*_ is the number of recipient phenotypes that phenotype *i* can reach by a single mutation of the mutated parameter under consideration.

The *phenotype-specific* mutation rate is proportional to the *general* mutation rate, represented by the parameter *m*. There is a great deal of variation in *m* among species and ecological contexts (Westra et al., 2017). In humans, mutations/base pair is estimated at ∼ 10^−8^ per generation (Nachman & Crowell, 2000) and, assuming an average gene size of ∼ 1000 base pairs, this results in a general mutation rate *m* on the order of 10^−5^ mutations/locus per generation. In *E. coli*, mutations/base pair is estimated at ∼10^−10^ per generation (Foster, et al., 2015). Thus, for an average gene size, the estimated general mutation rate *m* is on the order of 10^−7^ mutations/locus per generation. Matic et al. (1997) found that values for the *E. coli* mutation rate to drug resistance are typically in agreement with this figure (∼ 10^−7^), but they also found examples as high as ∼ 10^−5^. The values that might have been relevant for early periods of evolution are unknown, but likely to be on the higher end because of error-prone conditions thought to have prevailed at that time. This would be a relevant issue for our case study of a putative primordial circadian clock, which will be treated in **RESULTS**. We focus on spontaneous point mutations resulting from replication that are the major source of variation in a bacterium like *E. coli* (Foster, et al., 2015). The general mutation rate is subject to evolution in various contexts (Sniegowski et al., 2000; Raynes et al., 2018) and, as we will show for our clock model, the interaction of phenotype-specific mutation rates and near-neutral fitness effects (i.e., growth rates, as a measure of total fitness, are nearly equal for all phenotypes) results in different values of the general mutation rate that are optimal for individual phenotypes.

Finally, the *phenotype*-*specific* mutation rate constant *m*_*ij*_ between phenotypes *i* and *j* is given by the product of two factors *mk*_*ij*_, where *m* is the *general* mutation rate constant given by the number of mutations per generation and *k*_*ij*_ is the probability of a specific transition from phenotype *i* to phenotype *j*. The production rate of a phenotype (mutant) in units of mutations/time is then the product of *m*_*ij*_ the phenotype-specific mutation rate constant, *γ* _*i*_ the *exponential growth rate constant* of the donor phenotype (related to the doubling time, ln 2 / *τ* _*D*_), and *N*_*i*_ the size of the donor population. Note that this differs from the conventional description in that the product *mk*_*ij*_ is typically represented by a single *specific* rate constant per generation (e.g., Levin et al., 2000; Reams et al., 2010) that is not predicted but measured or estimated for a particular mutant phenotype.

### Population Dynamic Equations

To simplify the presentation, we restrict consideration to asexual haploid organisms in a spatially homogenous context growing in an exponential steady state, which is the most rigorously defined state for a cellular population (Maaløe & Kjeldgaard, 1966). Under these idealized conditions, all effective population sizes *N*_*e*_ are equal to the census population size *N*, mutants are never lost from the population, and the equilibrium distribution can be rigorously determined under non-selecting and selecting conditions. Lethal mutations (∼ 1%) can be subsumed within a net growth rate constant since there is evidence that these mutants occur by a first-order process (Robert et al., 2018). Of course, steady-state exponential growth cannot continue indefinitely. Nevertheless, results obtained under these conditions provide a rigorous reference or standard to which results under more realistic conditions can be compared, analogous to the historical role played by the frictionless plane in mechanics (Hawking, 2002) and by the Hardy-Weinberg law in population genetics (Crow, 1988; Wakeley, 2005). As with these idealizations, the intention is to get at something essential with the understanding that refinements will undoubtedly be added in the future; just as wind resistance and static friction were eventually added in mechanics and selection, drift and population structure were eventually added in population genetics. In each case, the expectation is that more realistic aspects will be added as the theory becomes refined. In **DISCUSSION** we will suggest methods to relax our assumptions.

The population dynamic equations for steady-state exponential growth can be written in terms of numbers for each of the *n* phenotypes in the population

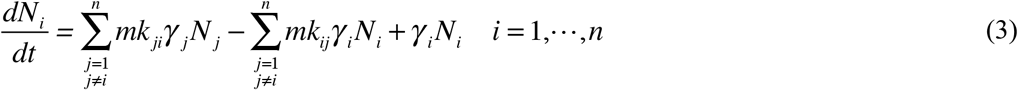

The first sum is the rate of increase by mutation, the second sum the rate of loss by mutation, and the final term the rate of increase by net exponential growth, with *γ* _*i*_ in doublings per unit time. These equations have the undesirable feature that the population is continually increasing. However, by expressing the population numbers *N*_*i*_ as a fraction of the total population *N*_*T*_ (or relative frequency) the resulting equations have a more convenient form with a well-defined steady state. Thus, the relative frequency of phenotype *i* is

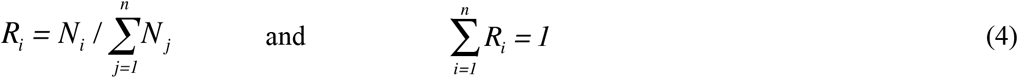

Starting with the derivative of the relative frequency

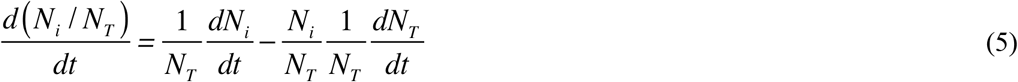

substituting *dN*_*i*_ */ dt* from **Eqn. (3)**, and noting the cancelation of the mutation terms in *dN*_*T*_ */ dt*, we obtain

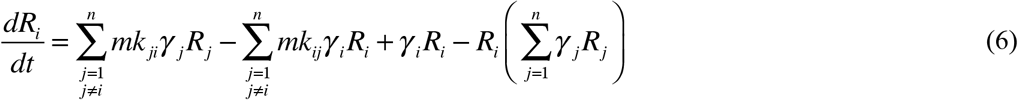

In anticipation of the case study to follow, we shall consider the situation in which phenotype *k* has growth rate *γ* _*k*_ in a *non-selecting condition* and 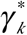 in a *selecting condition*. By adding and subtracting the same terms, normalizing time *t* by *γ* _*k*_ (*τ* = *γ* _*k*_ *t* in generations) and defining relative growth rates as *μ*_*i*_ = *γ* _*i*_ */ γ* _*k*_, **Eqn. (6)** can be rearranged and rewritten for phenotype *k* and for all other phenotypes *i* to emphasize three separate contributions to their rate of change:

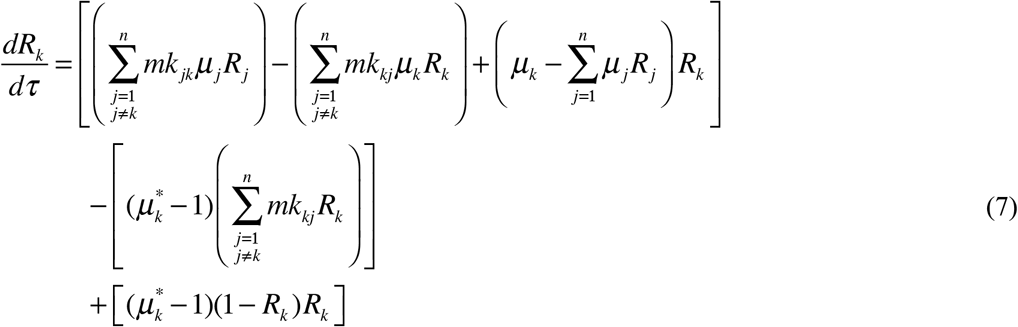

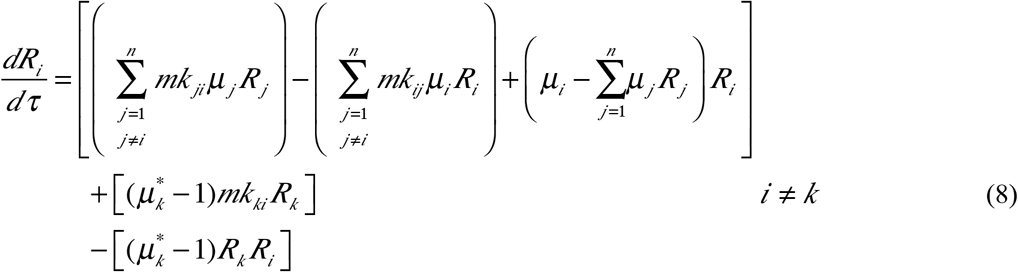

where the seledtion coefficient is defined as 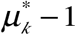. If there are no fitness effects in the *non-selecting* condition (all growth rates identical), then the form of the above equations in the *selecting* condition has the meaning

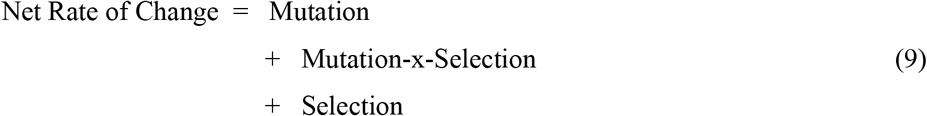

The middle term involves mutations generated specifically by replication of the phenotype with the selective advantage; hence, it is the only term that involves both a mutation rate and the selection coefficient. The above equations can be considered one of several alternative forms of the standard population genetic equations (Wilke, 2005); however, the alternative form used here most clearly reveals the three distinct rate contributions we wish to consider.

## RESULTS

The Design Space Toolbox 3.0 has algorithms for the automatic prediction of numerous characteristics within and between phenotypes. Examples of characteristics *within* phenotypes include the predicted volume (global robustness) of individual phenotypes, protein burden due to differential protein expression, dynamic behavior, and system design principles for the realization of the phenotype. Volumes are shown with identifying phenotype numbers in **Figure 2A**. There are numerous phenotypic characteristics that can be plotted on the z-axis as a heat map; an example is the protein burden of each phenotype due to differential protein expression in the non-selecting condition (**Figure 2B**). Simulation of the full system, with time *t* scaled by a factor of 1/3 (*τ* = *t / 3*) to match a 24-hour cycle time, produces the results in **Figure 2C**, which validates the prediction of a damped oscillatory characteristic for phenotype #7.

Examples of characteristics that distinguish *between* phenotypes include phenotype-specific mutation rate constants and system design principles. The first example, involving phenotype-specific mutation rate constants, distinguish between phenotypes in the context of the population dynamic model. In this model the focus is on the dynamics of the populations of organisms with the different phenotypes rather than the dynamics of the biochemical molecules of the system (oscillator). The second example, involving design principles, follows from the definition of a “qualitatively-distinct phenotype” as a combination of dominant processes operating within an intact biochemical system (Savageau et al, 2009). Phenotype #5 (signature 11 11 21 11 **11** 11) and phenotype #7 (signature 11 11 21 11 **21** 11) in **Table 1** are distinguished by a single change in dominance involving the rate of transcription of the mRNA for the activator (Bold digits in the signature). In our simplified case, allowing only the two equilibrium dissociation constants to vary by mutation, the condition is the following: Phenotype #5 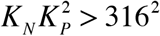 and Phenotype #7 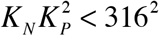. This suggests that a mutation increasing *K*_*N*_ alone by a sufficient amount can convert phenotype #7 to #5. In the more general context of distinguishing phenotype #7 from its neighbors, phenotype #7 in **Figure 2A** must be to the left of phenotypes #5 and #8 and to the right of phenotypes #3 and #15. The result is not at all obvious or intuitive, rather it is a subtle *system design principle* (Savageau & Fasini, 2009; Savageau, 2013) defined by four boundaries (**Supplemental Information, Section S10**). Thus, all system parameters must satisfy constraints involving specific constellations of values with many opportunities for compensation; there is no single parameter capable of distinguishing between phenotypes. This is particularly apparent in the case of complex diseases for which many genes and parameters interact in subtle ways that are difficult to identify; there is no single effective target for treatment, rather there are many potential targets with a spectrum of effectiveness.

Small changes, in the limit of linearization, within a phenotypic region eliminates the possibility of epistatic interactions. Larger changes, but still within a phenotypic region, can account for a variety of epistatic interactions. For example, the simple conditions in the previous paragraph show an epistatic interaction between two mutations with one affecting *K*_*N*_ and the other affecting *K*_*P*_. This is clear from the fundamental product of power law nonlinearities found in biochemical kinetics. Moreover, with changes large enough to move the system from one qualitatively distinct phenotypic region to another, nearly any type of epistatic interaction can be realized.

### Fixing Two Free Parameters

Two features that any population model should capture are that “large-effect” mutations are rare whereas “small-effect” mutations are abundant in well adapted systems (Bataillon & Bailey, 2014; Tataru et al., 2017; Bondel et al., 2019; Templeton, 2021) and detrimental mutations outnumber beneficial ones. Although there are exceptions, which we discuss later, these two features must be considered in the context of our model before we can predict phenotype-specific mutation rate constants and fitness effects.

Although terms such as large-, small-, zero-, positive-, and negative-effect are often applied to mutations in describing their effects on *fitness*, these terms only apply to populations in a given environment. With a change in environment the same mutation can have a different, indeed often an opposite, effect on fitness (Templeton, 2021). This is because fitness is a property of the phenotype, which in turn is a function of both genotype and environment. To separate these issues, we use the terms “size scale” (i.e., whether the change in value of a kinetic parameter caused by mutation is large or small) and “directional bias” (i.e., whether parameter change caused by mutation is in the direction of increasing or decreasing entropy) to characterize mutations without regard to fitness. Fitness is then a function of the phenotype and not of the mutation per se. This separation has the advantage of allowing us to characterize the frequency distribution of phenotypes under non-selecting and selecting conditions.

In our theoretical framework, we account for the size scale and directional bias of mutations with an exponential distribution having scale factor λ and directional bias parameter δ that increases or decreases the effective scale factor. Unlike the other parameters in our theoretical framework, these two must be estimated from experimental data. For this purpose, we draw upon the best studied specific function in molecular biology, LAC repressor binding to its recognition sites in the DNA of *E. coli*. Markiewicz et al. (1994) generated ∼4000 protein variants by making substitutions at each amino acid position. After being transformed through the molecular mechanisms that provide the causal connection between the gene sequence and the integrated function of the *lac* system, a corresponding distribution of phenotypes was determined. As Markiewicz et al. (1994) showed, there are essentially three qualitatively-distinct phenotypes involving LAC binding: (1) the inducible “wild-type”, (2) non-inducible constitutive, and (3) non-inducible super-repressed. Under the conventional laboratory conditions used to detect these three phenotypes, the data in their figure 1 show that changes at ∼ 67% of the positions were tolerant to substitutions (no change in DNA binding), 31% were intolerant with an increase in binding entropy (decrease in DNA binding), and ∼2% were intolerant with a decrease in entropy (increase in DNA binding).

In our prediction of phenotypes resulting from mutations in the N gene of the clock model, there are three analogous qualitatively-distinct phenotypes: the oscillatory “wild type”, the non-oscillatory constitutive, and non-oscillatory super-repressed (**Figure S6**). Values of λ=0.6 and δ=1.85 provide the best fit to the experimental data and the predicted distribution of fitness effects in this case is ∼ 67% oscillatory (wild-type DNA binding), ∼ 31% non-oscillatory constitutive (decreased DNA binding), and ∼ 2% non-oscillatory super-repressed (increased DNA binding). This distribution of effects is likely to be similar for other proteins at least qualitatively. For example, Markiewicz et al. (1994) examined the sequence alignment of proteins in the LAC family of proteins (which includes proteins of unrelated function in addition to other transcription factors) and found that 61% of the residues were not conserved (tolerant of evolutionary changes) and 39% were conserved (intolerant of evolutionary changes). In view of the different percentages of non-conserved and conserved residues being similar in a variety of unrelated proteins, fitting values of the λ and δ to these percentages is likely to be appropriate for other systems as well.

To summarize, there are two free parameters in this model, λ and δ, that must be estimated from experimental data. Based on the above considerations, we assign the following model values for these two parameters: λ=0.6 and δ=1.85. All the remaining parameters have values predicted solely based on the underlying mechanistic model using methods from the Design Space Toolbox (Valderrama-Gómez et al, 2020), as described in the following sections.

### Phenotype-Specific Mutation Rate Constants

In what follows we predict the equilibrium distribution of phenotype diversity under non-selecting conditions in three stages to clearly distinguish different contributions. First, we consider the idealized case in which there is no size scale or directional bias for mutations that have neutral fitness effects and show that the distribution differs from the expectation of a uniform distribution. Second, we add size scale and directional bias and find that the distribution exhibits an increasing gradient from phenotypes with low entropy to those with high entropy. Third, as a specific example involving phenotypes with mixed fitness effects, we consider their protein burdens to obtain a distribution with a central peak resulting from *entropy – selection balance*. It should be noted that this type of balance is different from other types of specific mutation – selection balance (Barton, 2007; Lynch, 2010; Orlenko et al., 2016) and the general mutation – selection balance that always exists at equilibrium. Finally, we illustrate the shift in the distribution when the oscillatory phenotype is subject to various degrees of selection.

### Distributions for Neutral Mutations Without Size Scale or Directional Bias Effects

Neutral mutations without size scale or directional bias effects produce a uniform distribution of values in parameter space; however, the partitioning of design space, which is dictated by the architecture of the underlying molecular system, results in an equilibrium distribution of phenotype frequencies that is determined by the normalized values of the phenotypic volumes (**Figure 3A**, Blue), as obtained analytically. Large volumes (e.g., phenotype #6) imply robust phenotypes that are tolerant to large changes in parameter values; small volumes (e.g., phenotype #15) imply fragile phenotypes that are easily disrupted by small changes. The absence of size scale and directional bias is of course an idealization, but useful for identifying the volume contribution and providing a baseline on which to characterize more realistic features, as described below.

**Figure 3.**
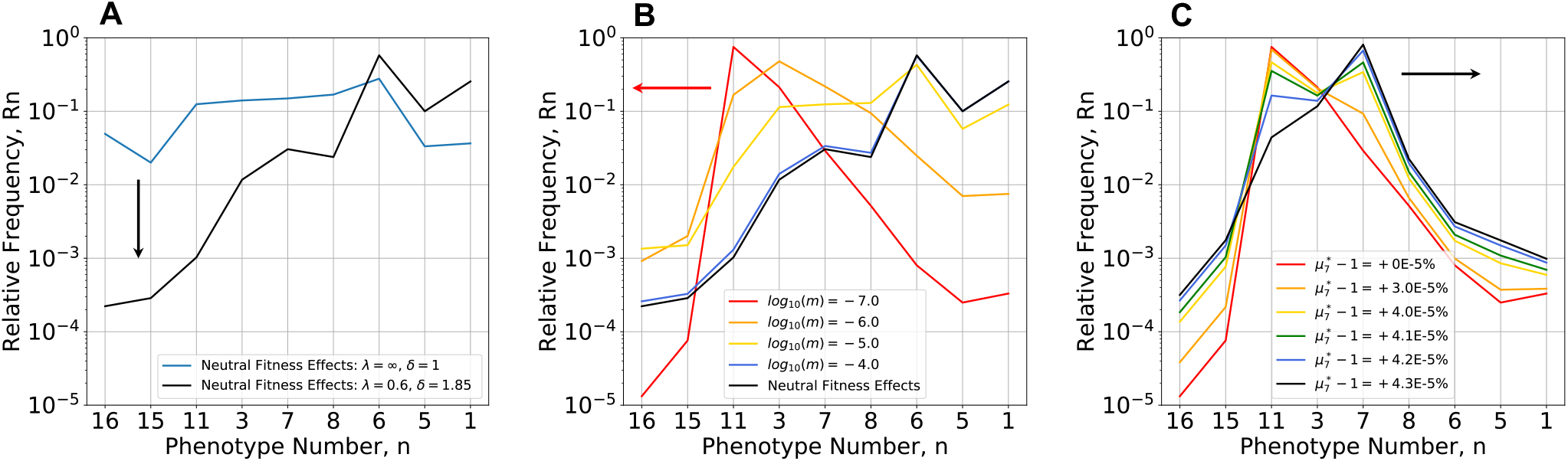
Equilibrium distributions of phenotype diversity. Mutational entropy is increasing from left to right, from the phenotype with both equilibrium dissociation constants having the lowest values (phenotype #16) to that with both having the highest values (phenotype #1). **(A) Mutations with neutral fitness effects (all** *μ*_*i*_ = 1) **under non-selecting conditions** (Blue) in the absence of size scale (*λ* → *∞*) and directional bias (***δ*** = 1), and shifted down (Black) in the presence of size scale (*λ* = 0.6) and directional bias (***δ*** = 1.85). In the absence of directional bias there is a minimal gradient; whereas this gradient is approximately 4-orders of magnitude when directional bias is present. **(B) Mutations with mixed fitness effects** (*μ*_*i*_ different) **under non-selecting conditions** in the presence of size scale (*λ* = 0.6) and directional bias (***δ*** = 1.85). The distribution is shifted to the left with decreasing values of m =10^−4^ (Blue), 10^−5^ (Yellow), 10^−6^ (Orange) and 10^−7^ (Red) compared with the strictly neutral results in (Black). The distribution changes dramatically, increasing, reaching a peak, and then decreasing when directional bias is present. Fitness effects normalized with respect to the experimental data for *E. coli* ß-galactosidase burden. **(C) Mutations with mixed fitness effects** (*μ*_*i*_ **different**) **under selecting conditions with various degrees of selection**. The peak of the distribution under the non-selecting conditions (*m*=10^−7^) shifts to the right, from phenotype #11 (non-oscillatory, Red) to phenotype #7 (oscillatory, Black) and its frequency increases with increasing values of the selection coefficient whereas the frequency of the other phenotypes decrease according to their selective disadvantage.

### Distributions for Neutral Mutations with Size Scale and Directional Bias Effects

In the presence of size scale and directional bias effects (λ = 0.6 and δ = 1.85), the equilibrium distribution exhibits a gradient from phenotypes with lower entropy (lower left corner in **Figure 2A**) toward phenotypes with higher entropy (upper right corner in **Figure 2A**), as obtained numerically from the steady state solution of the population dynamic equations and shown in **Figure 3A (**Black). Note that the phenotype with highest entropy, based on directional bias, is phenotype #1, which corresponds to mutations in both transcription factors that essentially eliminates the ability to recognize their DNA binding sites. Conversely, the phenotype with the lowest entropy, based on directional bias, is phenotype #16, which corresponds to mutations in both transcription factors that makes for overly tight binding. The gradient in this case is approximately 4-orders of magnitude.

### Distributions for Mixed Mutations with Size Scale and Directional Bias Effects

In the non-selecting constant light environment, in which mutants are assumed to exhibit fitness differences unrelated to the specific phenotype characteristic of oscillation, the equilibrium distribution is among mutations with mixed fitness effects, positive, negative and neutral. As an example of a phenotype-specific fitness characteristic that can be predicted, we consider the size of the protein coding regions and the protein burden of extraneous protein expression for each phenotype.

Experimental evidence in the case of *lac* operon expression in *E. coli* suggests that inappropriate constitutive expression (nevertheless within the normal range for expression of the wild-type induced state) decreases the growth rate by < ∼ 0.1%. (Koch, 1983). The decrease would be even less if we consider only the contribution from ß-galactosidase, and neglecting that from the permease and transacetylase, in making estimates for our clock module. Given the 10-fold larger size of the ß-galactosidase monomer, its tetrameric structure and the 1000-fold protein burden (difference between wild-type uninduced expressed and mutant constitutive expression), compared to the assumed 100 amino acid length, dimer structure and predicted 100-fold protein burden for our molecular model, allows the appropriately scaled decrease in growth rate to be < ∼ 0.001%. The following relative growth rates (selection coefficients) for each phenotype, relative to phenotype #7 in the non-selecting condition, follow from the predicted levels of protein expression for each phenotype (**Figure 3B**): *μ*_1_ = 0.999997573 (−2.43E-04 %), *μ*_3_ = 1.000000322 (3.22E-05 %), *μ*_5_ = 0.999997527 (−2.47E-04 %), *μ*_6_ = 0.999997357 (−2.64E-04 %), *μ*_7_ = 1.0 (0 %), *μ*_8_ = 0.999999693 (- 3.07E-05 %), *μ*_11_ = 1.000000412 (4.12E-05 %), *μ*_15_ = 1.000000115 (1.15E-05 %), and *μ*_16_ = 0.99999976 (−2.40E-05 %). Note that these small differences in growth rate that are undoubtedly overestimates would be considered neutral, given the technical limitations of experimentally determining growth rate differences less than ∼ 0.1% (Gallet et al., 2012).

When both size scale and directional bias effects are present, the graded distribution in the strictly neutral case (**Figure 3A,B**, Black) is dramatically changed to a peaked distribution that is increasingly weighted to the left (**Figure 3B** Orange, Red) as the general mutation rate is decreased. The result is what might be called *entropy-selection balance*.

Note that all the distributions in **Figure 3A** and **3B** occur under the *non-selecting condition with respect to the oscillatory* phenotype characteristic. Moreover, despite the difficulty distinguishing between mutations with neutral fitness effects and mutations without detectable fitness effects, these results show that the equilibrium distributions are radically different. It is also clear that there is an optimal value for the general mutation rate that favors each phenotype.

### Equilibrium Distribution of Phenotype Diversity Under the Selecting Condition

When connections to both the synchronizing environmental signal and the integrated cellular biochemistry are made by a critical new mutation, it would confer no selective advantage if it were to occur in one of the phenotypic regions that lack the ability to oscillate. For example, it has the highest probability of occurring in phenotype #11 because its frequency in the population is nearly 100% before the mutation occurred. More rarely, it would occur in the region of phenotype #7, but then there would be the potential to synchronize with the light-dark environment (the selecting condition) and have a selective advantage. The predicted equilibrium distribution of phenotype diversity under the selecting condition as a function of the selection strength is shown in **Figure 3C**. Beyond a critical level of selection, the peak of the equilibrium distribution shifts from phenotype #11 to phenotype #7. Although we cannot currently predict the fitness of phenotype #7 under selecting conditions, if it were possible to estimate the distribution of phenotype diversity, then one could back calculate the selection strength that produces the best fit to the estimated distribution (**Supplemental Information, Section S11**).

The three separate contributions to the rate of change in phenotype frequency in the neutral case (**Eqn. 9**, mutation, mutation-x-selection, and selection) are shown in **Figure 4** as a function of selection strength and general mutation rate. The rate of change at equilibrium is equal to zero and the contributions of mutation alone and selection alone are nearly opposite and equal. The contribution from mutation-x-selection is negligible at the selection strengths shown. Note the differences in scale: the maximum contribution to the rate at equilibrium is proportional to the general mutation rate, and the degree of selection necessary to achieve the maximum rate increases rapidly with the general mutation rate.

**Figure 4.**
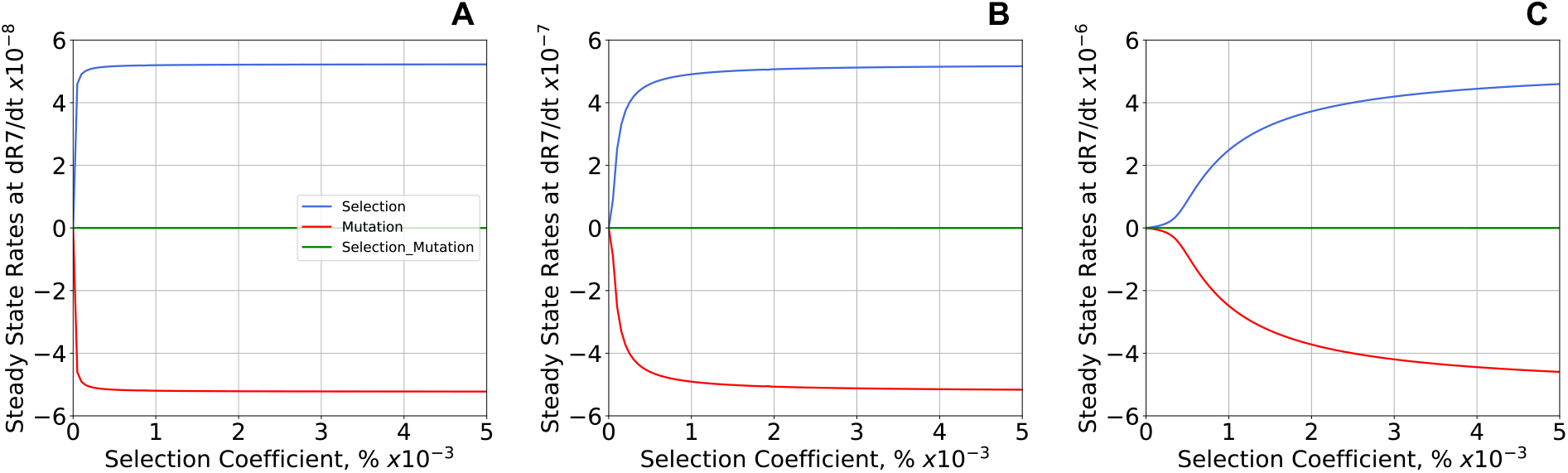
Three separate contributions to the steady-state rate of change in frequency for the oscillatory phenotype #7. The three panels show results for mutations with neutral fitness effects and general mutation rate *m* = 10^−7^ (**A**), *m* = 10^−6^ (**B**), and *m* = 10^−5^ (**C**). The contributions (**Eqn. 9**) are shown as a function of selection strength at equilibrium. Selection alone (Blue) is balanced with mutation alone (Red); the contribution by mutation-x-selection (Green) is negligible for the strengths of selection shown. The maximum rates of change are proportional to the general mutation rate (note the change of scales), and stronger selection is required to overcome the effects of higher general mutation rates.

### Non-Equilibrium Distribution of Phenotypes Under the Selecting Condition

In this and the following subsection, instead of determining the phenotype distribution at equilibrium under either the non-selecting or selecting condition, we determine the temporal changes in distribution between the two equilibria – from non-selecting to selecting or from selection to non-selecting. The light-dark environment generates the selecting condition. The ability to synchronize with the light-dark environment generates a selective advantage for the oscillatory phenotype (#7) greater than that of the other phenotypes. Aside from #7, all the other phenotypes have either a mixed distribution or a neutral distribution of fitness effects.

Results with a neutral distribution of fitness effects for phenotypes other than #7 are shown in **Figure 5A**, starting from the equilibrium distribution under the non-selecting condition (**Figure 3A,B**: Black) and evolving to the equilibrium distribution under the selecting condition (selection coefficient μ_7_^*^ - 1 = 6.0E-3 %, all other μ_*i*_ = μ_7_ and fixed). Phenotype #7 increases rapidly with a time scale dominated by selection, while there is little change in the other phenotypes until ∼ 5.0E+04 generations (**Figure 5A**, vertical dashed line). After this point, phenotype #7 approaches its maximum at ∼ 1.5E+05 and all other phenotypes slowly decrease asymptotically toward the new equilibrium distribution with a time scale dominated by mutation. There are no changes in the ranking of phenotype frequencies in the population after 3.5E+05 generations.

**Figure 5.**
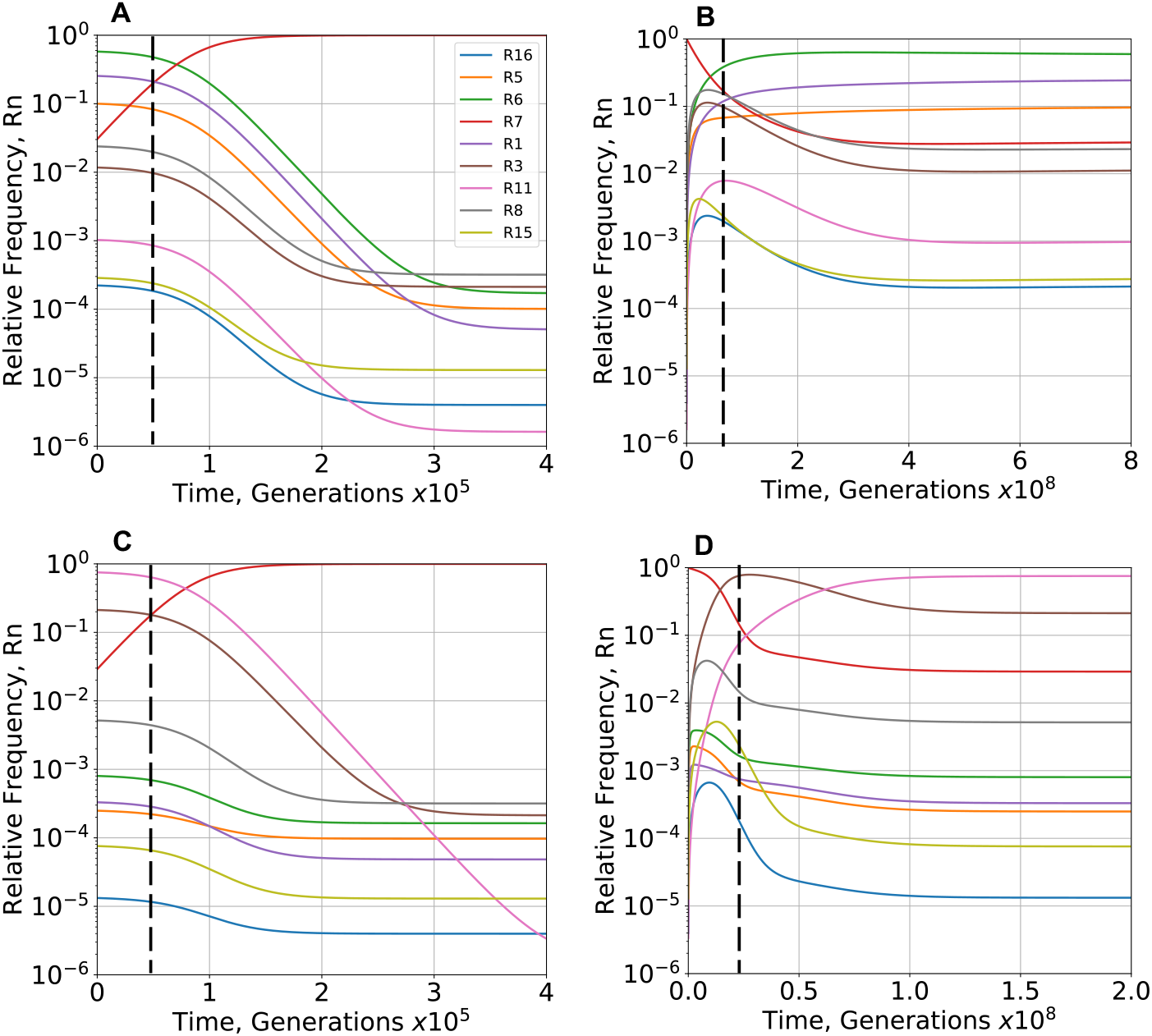
Temporal response in relative frequency of phenotypes following imposition and removal of the selecting condition. **(A,B)** Neutral distribution of fitness effects. **(A)** The increase of phenotype #7 (Red) is accompanied initially by very little change in the other phenotypes, followed (after the dashed line) by a slow decrease in all other phenotypes. All changes in the rank of the relative frequencies occur before 3.5E+05 generations. **(B)** The decrease of phenotype #7 (Red) is accompanied initially by a rapid increase in all other phenotypes, a peak (the last occurring at the dashed line) followed by a slow decrease in all other phenotypes except for phenotype #1, #5 and #6, which continue to increase slowly. All changes in the rank of the relative frequencies occur before 2.5E+08 generations. The overall response is ∼ 1000-times slower than **(A). (C**,**D)** Mixed distribution of fitness effects. **(C)** The increase of phenotype #7 (Red) is accompanied initially by very little change in the other phenotypes, followed (after the dashed line) by a slow decrease in all other phenotypes. All changes in the rank of the relative frequencies occur before 4.0E+05 generations. **(D)** The decrease of phenotype #7 (Red) is accompanied initially by a rapid increase in all other phenotypes, a peak (the last occurring at the dashed line) followed by a slow decrease in all other phenotypes except for phenotype #11, which continues to increase. All changes in the rank of the relative frequencies occur before 6.3E+07 generations. The overall response is ∼ 400-times slower than **(C)**. Imposition occurs by a change from a non-selecting (μ_7_* = 1.0) to a selecting (μ_7_* = 1.00006) environment and removal by the reverse. The general mutation rate *m* = 10^−7^.

### Non-Equilibrium Distribution of Phenotypes with Removal of the Selecting Condition

Experimental studies have explored the evolutionary loss of phenotypes in response to the relaxation of selection. For example, the ability of *Bacillus subtilis* to sporulate is lost when it is no longer under selection (Maughan et al., 2007). In the clock model, relaxation of selection occurs when the selective advantage of phenotype #7 is removed (switched to constant light) and the population returns with time to the equilibrium distribution under the non-selecting condition.

Results with a neutral distribution of fitness effects for phenotypes other than #7 are shown in **Figure 5B**, starting from the equilibrium distribution under the selecting condition (selection coefficient μ_7_^*^ - 1 = 6.0E-3 %, all other μ_*i*_ = μ_7_ and fixed) and evolving to the distribution under the non-selecting condition (**Figure 3A**,**B**: Black). The large number of the selected phenotype (#7) in the initial equilibrium distribution is rapidly lost and redistributed to all the other phenotypes within ∼ 7.5E+07 generations. There is a subsequent slow redistribution and decrease among all the phenotypes except #1, #5 and #6 (high entropy phenotypes) until a new equilibrium distribution is approached asymptotically with a time scale dominated by mutation. There are no changes in the ranking of phenotype frequencies in the population after ∼ 2.5E+08 generations. Comparison of the time scales in **Figure 5A and B** shows that the response to the removal of selection is approximately ∼ 1000-times slower than that to the imposition of selection.

Results with a mixed distribution of fitness effects for phenotypes other than #7 are shown in **Figure 5D**, starting from the equilibrium distribution under the selecting condition (selection coefficient μ_7_^*^ - 1 = 6.0E-3 %, all other μ_*i*_ determined by protein burden and fixed) and evolving to the distribution under the non-selecting condition (**Figure 3B**: red). The large number of the selected phenotype (#7) in the initial equilibrium distribution is rapidly lost and redistributed to all the other phenotypes within ∼ 2.5E+07 generations. There is a subsequent slow redistribution and decrease among all the phenotypes except #11 (low entropy phenotype) until a new equilibrium distribution is approached asymptotically with a time scale dominated by mutation. There are no changes in the ranking of phenotype frequencies in the population after ∼ 6.3E+07 generations. These results are in qualitative agreement with those of Maughan et al. (2007) when the larger target size of the sporulation machinery and the higher mutation rate of their mutator strain are considered. Comparison of the time scales in **Figure 5C and D** shows that the response to the removal of selection is approximately ∼ 400-times slower than that to the imposition of selection.

The large differences in time scale indicate that alternating between equal periods in selecting and non-selecting environments before reaching equilibria would lead not to an average of the two distributions but to a distribution closer to that in the selecting environment, which is reminiscent of “conflict between selection in two directions” (Haldane & Jayakar, 1963).

## EXPERIMENTAL IMPLICATIONS

There are two major challenges in determining the distribution of phenotypes available for selection to act upon. One is the time of sampling relative to the evolutionary dynamics of natural populations and the second is technical limitations in the ability to identify and measure phenotypes. Both help to explain the pessimism expressed by Charlesworth (1996) in determining the distribution of the phenotypes and their fitness characteristics in natural populations.

Experimental studies based on mutants constructed from a highly evolved system (wild type) in a given environment (in the extreme, optimized according to Fisher’s Geometric model) may have only a very narrow distribution of alternative phenotypes capable of improvement in that environment. Those based on mutants constructed from a system that is far from its optimal state in a new environment, are likely to offer a more fertile distribution of phenotypes capable of improvement. Indeed, Matuszewski et al. (2014) pointed out a violation of Fisher’s prediction that mutations of small effect are the primary raw material of adaptive evolution. They considered a geometric model like Fisher’s but with environmental change. In contrast to Fisher’s predictions, larger adaptive steps often occur with a moving optimum. Mutations of small effect are not always the main material of adaptive change even when there is a single adaptive optimum, albeit a moving one. However, determining the natural distribution from subsequent measurements depends on the time of sampling following the construction, with the actual distribution of fitness effects bounded by two extremes: sampling at time zero and sampling at the time to reach equilibrium. The time zero sample has not involved any exchange; thus, it simply reflects the construction and may have little to do with any subsequent distribution in nature. The equilibrium sample in some cases might be the more relevant distribution in nature, but there is insufficient time to test this in practice. Thus, the natural distribution undoubtedly lies somewhere between these extremes. There is the additional difficulty of identifying the phenotypes because of technical limitations. Experimental studies based on natural variants face the same two challenges.

Orr (2005) also identifies challenges in two related problems. “The first is the current theory is limited in several ways – all the models that have been mentioned rest on important assumptions and idealizations. Although they are reasonable starting points for theory, none of these assumptions is necessarily correct and changing any might well change our predictions. […] The second problem concerns testability. The difficulty is practical, not principled. Whereas current theory does make testable predictions, the effort required to perform these tests is often enormous (particularly as the theory is probabilistic, making predictions over many realizations of adaptation). Given, for example, the inevitable and often severe limits on replication in microbial evolution work, we can usually do no more than test qualitative predictions.” Our theory is grounded in measurable biochemical parameters, and thus a different set of assumptions and idealizations need experimental testing.

### Experimental Evolution Studies in a Chemostat

The equilibrium distributions of phenotype diversity under selecting and non-selecting conditions can be approximated experimentally by growing populations in a chemostat/turbidostat (Bustos & Golden, 1992; Gresham & Jong, 2015). This allows us to relax the assumptions concerning the ideal context. If a one-liter chemostat is initialized with a single cell and the population grows exponentially until reaching typical densities of 10^8^ to 10^10^ cells/ml (Gresham & Jong, 2015), at this point nearly all phenotypes will be present in the population (**Figure 6**). If the flow of fresh media into the chemostat is initiated at this point, the doubling of the population in each subsequent generation due to growth coupled with the 50% reduction in population size per generation due to dilution will introduce fluctuations in the numbers of cells. Phenotypes with a low frequency must be treated stochastically when the differences between effective population size and the census population size become significant. All other phenotypes are expected to persist in the chemostat.

**Figure 6.**
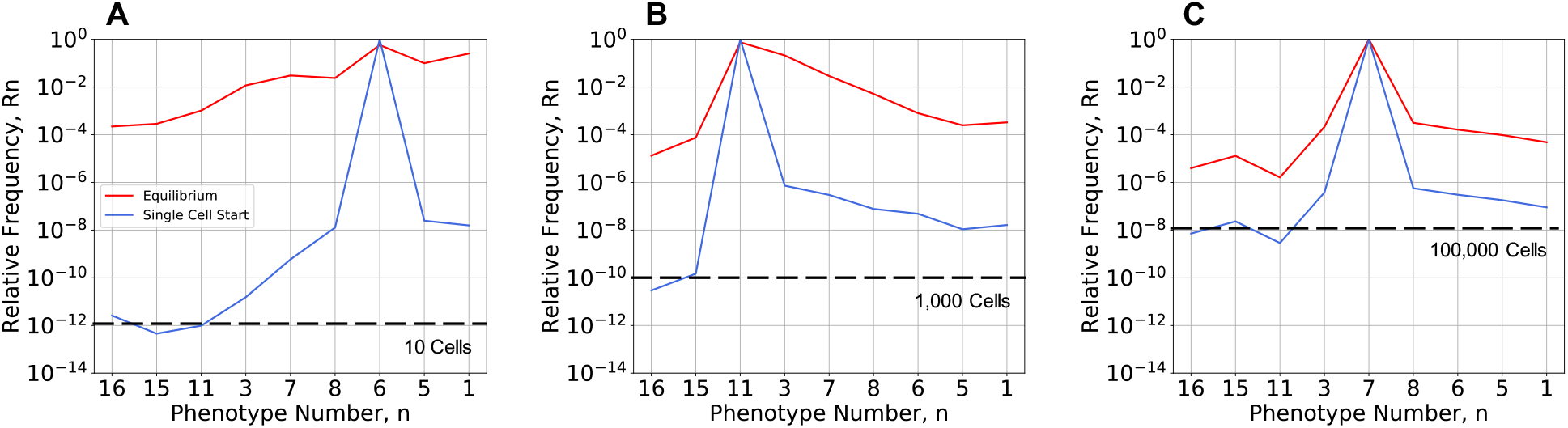
Non-equilibrium distributions of phenotype diversity under non-selecting and selecting conditions after exponential grow from one to 10^13^ cells. Cells with general mutation rate *m* = 10^−7^ are inoculated into fresh media in a one-liter chemostat without flow. **(A)** Under non-selecting conditions with neutral fitness effects, phenotypes with the lowest frequency (#11, #15 and #16) are expected to have ∼ 10 cells in the chemostat. **(B)** Under non-selecting conditions with a protein burden spectrum of fitness effects, phenotypes with the lowest frequency (#15 and #16) are expected to number ∼ 1000 cells. **(C)** Under selecting conditions with a protein burden spectrum of fitness effects, nearly all phenotypes are expected to be present at more than ∼ 100,000 cells. Size scale effects and directional bias effects are present in all cases. The initial distribution (Blue) can be expected to approach the equilibrium distribution (Red) asymptotically with time following long-term exponential growth with the flow of fresh media to the chemostat.

Under non-selecting conditions, the case with neutral fitness effects is the most difficult. At the time when the chemostat reaches the full operating density, all phenotypes in the population will be present with a significant frequency except for #11, #15 and #16 (∼ 10 cells in **Figure 6A**). The issue of genetic drift could be introduced here by the addition of stochastic changes (replication or removal) in each generation. In the case of mixed fitness effects due to protein burden, even those phenotypes with the smallest frequency will have a census size of ∼ 1000 cells (**Figure 6B**). Under selecting conditions, at the time when the chemostat reaches the full operating density, even those phenotypes with the smallest frequency will have a census size of ∼ 100,000 cells (**Figure 6C**).

### Measuring Qualitatively Distinct Phenotypes

Although current experimental limitations make it difficult to measure individual phenotypes, there are some cases in which relevant aggregate phenotypes can be measured. In the classic studies of Markiewicz et al. (1994), the authors constructed a collection of LAC mutants, measured their β-galactosidase expression, grouped the results into qualitatively-distinct phenotypes (constitutive, super-repressed or inducible), and determined the resulting distribution of phenotypes *measured at time zero*. They found 2% super-repressed, 67% inducible, and 31% constitutive. This is not surprising, given a low mutation rate (*m* = 10^−7^) and that the construction started with the highly evolved and presumably fit *lac* system of *E. coli*.

Fluorescently tagged protein might be an updated approach for other proteins. In our case study, measuring the activity of the N gene protein and classifying the results as constitutive (phenotypes #6 and #8), super-repressed (phenotypes #3 and #11) or oscillatory (phenotype #7) leads to the following predictions. In analogy with the LAC studies, and sampling the distribution at time zero, our results would match those of Markiewicz et al. (1994) because these values were used to fit the two free parameters of our model λ and δ. The distribution of phenotypes measured after reaching equilibrium under non-selecting conditions (loss of selection) with *neutral fitness effects* is predicted to be 2% super-repressed, 5% oscillatory and 93% constitutive (**Figure 7A**, 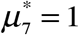). This reflects the dominant influence of entropy. The distribution of phenotypes measured after reaching equilibrium under non-selecting conditions with *mixed fitness effects* based on protein burden is predicted to be 96% super-repressed, 3% oscillatory and 1% constitutive (**Figure 7B**, 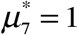). Under selecting conditions, the degree of selection required to reach a distribution with 60% oscillatory phenotype with mixed fitness effects is four-fold greater than that with neutral fitness effects. These differences, suggesting that the results with a neutral distribution of fitness effects can be achieved more easily than with the protein burden distribution, might be relevant for the evolution of LAC repressor as well. Furthermore, an examination of different values for the general mutation rate, *m*, at equilibrium with mixed fitness effects shows that even when the relative frequency of the oscillatory phenotype is maximum at *m* = 3×10^−6^, the results are still very different from that of wild-type LAC repressor selected in nature (**Supplemental Information, Section S11, Figure S7)**.

**Figure 7.**
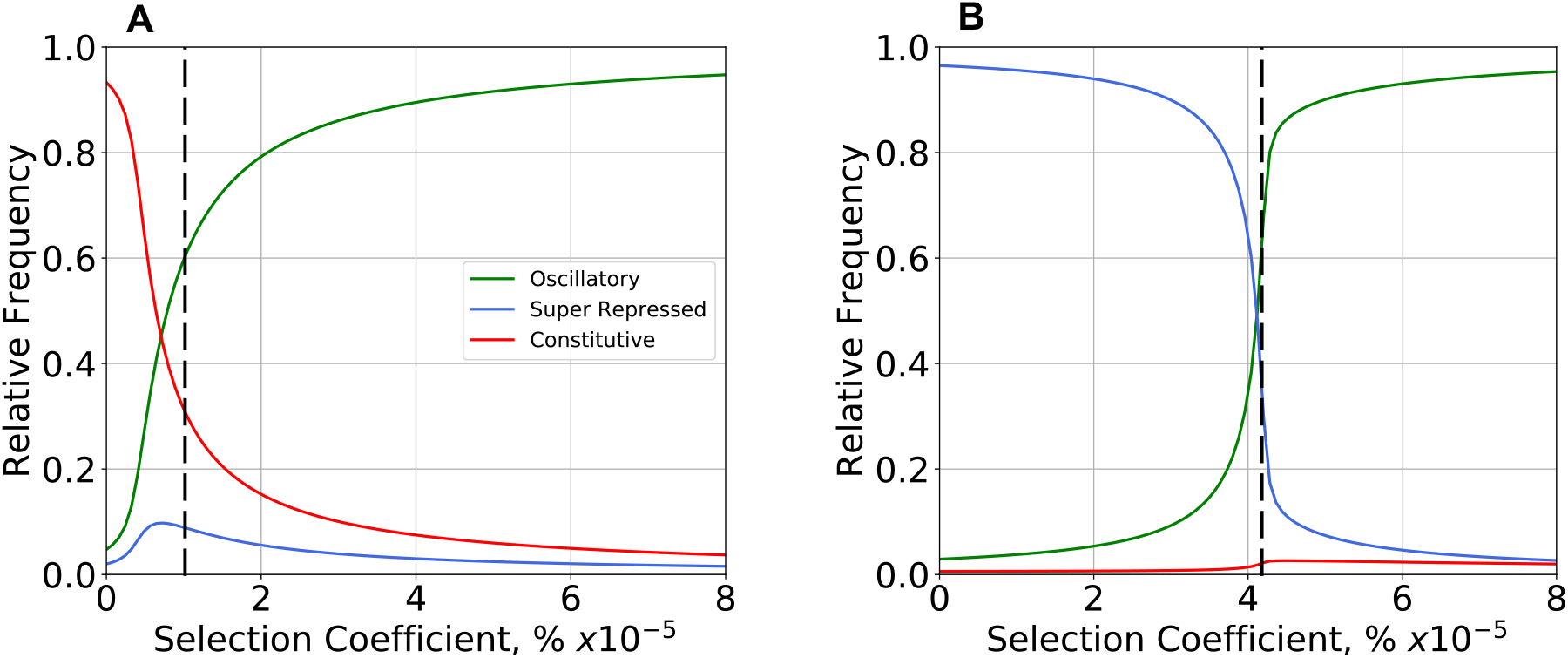
Equilibrium distributions of qualitatively distinct phenotypes under selecting conditions with various degrees of selection. Comparisons made with general mutation rate *m* = 10^−7^ and **(A)** neutral fitness effects (all *μ*_*i*_ = 1), and **(B)** a protein burden spectrum of fitness effects (*μ*_*i*_ different) under non-selecting conditions 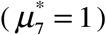. The degree of selection required to reach a distribution with 60% oscillatory phenotype (dashed line) with mixed fitness effects is ∼ four-fold greater than that with neutral fitness effects. The second most common phenotype is super-repressed with mixed and constitutive with neutral fitness effects. Thus, only the results predicted in in (**A**) match the experimental results of Markiewicz et al. (1994).

Testing such predictions would require finding rare cells in the population, at the limit of detection for many methods. Based on the start of a chemostat experiment as described in the previous section, the effluent at any subsequent time during the experimental evolution could be collected and the cells subjected to counting or sorting. Counting might well be able to determine the numbers of rare cells, sorting would allow sufficient material for further experimental tests. A double-sieve strategy would have advantages. First, the cells are grown under *non-selecting* conditions and sorted into two abundant classes, those with constitutive and non-constitutive expression. Second, the sorted cells with non-constitutive expression are grown under *selecting* conditions and sorted into those enriched for super-repressed and wild-type expression. This approach would require ∼ 10^10^ cells to be collected and sorted within a reasonable amount of time and cost, which should be feasible with recent advances in high-throughput sorting methods (Fan et al., 2013; Zhukov et al., 2021).

## DISCUSSION

Two complex and interrelated issues in evolution are the *distribution of phenotype diversity*, which offers opportunities for innovation, and the *interaction of phenotype-specific mutation rates and phenotype-fitness differences*, which determines population dynamics and the subsequent evolution of the population. Some experimental approaches to determining the distribution of mutant effects only address large effect mutations because there are technical limitations to the size of changes in growth rate that can be measured (Gallet et al., 2012). Others only address small effect mutations in the context of nearly-neutral theory (Kimura, 1983; Ohta, 1992). As Bondel, et al. (2019) pointed out, together the two provide a bigger picture by complementing one another. However, neither of these approaches deal with the causal linkages between genotype/environment and phenotype.

There are few examples attempting to determine the distribution of mutant effects by addressing the mechanistic link. Orlenko et al. (2016; 2017) have examined linear pathways in which classical Michaelis-Menten kinetics were assumed, kinetic parameters were sampled, and the system of ordinary differential equations was repeatedly solved. They found a lack of evolutionarily stable rate limiting steps in large stable populations (Orlenko et al., 2016) but stable patterns of limiting steps in some cases of fluctuating environments and population sizes (Orlenko et al., 2017). They note that more complex realistic systems remain to be studied in this context. Examples include systems involving more complex forms of regulation, enzyme-enzyme complexes and cascades, as well as branched and cyclic pathways. Loewe & Hillston (2008) focused on a simple limit cycle model for circadian rhythms (Leloup, et al., 1999) with a set of assumed parameter values as reference. They converted the biochemical kinetic equations from ordinary differential equations into pseudo-chemical kinetic equations for stochastic simulations. They employed dense sampling of parameter values and repeated stochastic simulations to generate statistical data for analysis in terms of various fitness correlates. Brajesh et al. (2019) focused on the *lac* operon of *E. coli* because it is a simple, specific system that has been studied for decades (Muller-Hill, 1996; Ullmann, 2003) and for which there are experimental values for nearly all the key parameters. Starting with this well-characterized system, they explored its phenotypic repertoire by dense sampling of the parameter space combined with numerical solution of the ordinary differential equations for the nonlinear mechanistic model. It will be difficult to replicate these approaches for other systems in which there is a large number of parameters with unknown value that are difficult or currently impossible to measure or estimate. This is precisely the bottleneck currently limiting the successful application of the conventional simulation-centric modeling strategy. This ultimately becomes a scaling issue for large systems because of the density of sampling required, coupled with the repeated deterministic and stochastic numerical simulations of the nonlinear differential equations. Moreover, with certain combinations of parameter values these numerical solutions often fail for technical reasons, which makes automation of the process problematic.

The phenotype-centric modeling strategy largely circumvents the bottleneck presented by a mechanistic model with a large number of unknown parameter values (Valderrama-Gómez & Savageau, 2018). Here we showed that it also can predict phenotype-specific mutation rates and the distribution of mutant effects under non-selecting and selecting conditions. It must be noted that the phenotype-centric approach does not escape the issue of scaling to large realistic systems, although it does not involve the limitations of dense sampling and repeated simulations mentioned above. The issue is the large number of phenotypes that must be treated analytically for any realistic system. However, each phenotype is a separate linear algebraic problem, which makes it what computer scientists call ‘embarrassingly parallelizable’, and therefore amenable to cloud computing.

By way of conclusion, we discuss differences between the theoretical framework of System Design Space and other theoretical frameworks, similarities between them, and potential areas of mutual interest for further development. We finish with a summary of results, some that are consistent with well-known results in theoretical population genetics and others that are new.

### Differences and Similarities Between Theoretical Frameworks

A broad context of theoretical population genetics is provided by the historical review of Orr (2005). He focused on the advances and limitations involving the two main classes of mathematical models: older phenotype-based models following in the spirit of Fisher’s geometric model and newer DNA sequence-based models emphasizing nearly neutral and extreme value theory.

The Design Space model has some superficial similarity to the geometric model of Fisher (1930), but it is fundamentally different. Although both prominently feature *geometry, quantitative phenotype traits* and *size of changes* caused by mutations, a brief comparison of Fisher’s Geometric model vs. the Design Space model shows there is little else in common:

- Phenotype definition is *generic, ad hoc, and descriptive* (height, weight, etc.) vs. *specific, mathematical, and rigorous* (genotypically-determined parameters and environmentally determined variables).
- Phenotypic traits are for *unspecified systems* in *unstructured Cartesian space* vs. *biochemically specified systems in structured logarithmic space*.
- Mutation causing *symmetric* changes involving *any combination of the orthogonal traits (omnidirectional)* vs. *asymmetric* (entropic) changes involving *one mechanistic trait (bidirectional)*.
- Mutations simultaneously affect *all n traits in general* (highly pleiotropic) vs. *n* = 1 *specific trait* (model-dependent pleiotropic).
- Organizing principle is *random variation in proximity to an optimum* vs. *deterministic structure of a Design Space*.
- Methodology focused on *statistical analysis and computer simulation* vs. *analytic geometry and computational algebra*.
- Focus on new mutations vs. standing genetic variation.

Although these theories are very different, there are a few connections between them that might be worth exploring. For example, two strong results from Fisher’s model and the extreme value theory are that an exponential distribution of positive effects mutations may be universal (Orr, 2005) and that there is a progression of size effects from initially large to subsequent smaller (Gillespie, 2004). In Design Space theory, the first of these results might have a connection to the assumption of exponential distributions for both positive and negative effect mutations. However, these exponential distributions are in logarithmic coordinates, which in design space theory means that they could also be considered power law in Cartesian coordinates. Regarding the second of the above results, we can speculate that if the initial mutation takes the system from an optimal state into a qualitatively different region of Design Space, then the first significant mutation taking it back will likely have a large effect on average. Once back near the optimum, then smaller quantitative changes will add refinements. However, back mutations with small changes at the level of kinetic parameters could lead to large qualitative changes at the phenotype level, but only when the phenotype undergoing back mutation is quantitatively near the common boundary with the recipient phenotype. This is also related to the long-standing robustness vs. evolvability issue (de Visser et al., 2003; Draghi et al., 2010; Payne & Wagner, 2014; Greenbury et al., 2016; Wei & Zhang, 2017). In our mechanistic framework phenotypes with large volumes in System Design Space are globally the most robust to mutation (to changes in the qualitatively distinct phenotype). Mutations with large-size effects can explore distant phenotypes infrequently. However, if there is a more favorable adjacent phenotype, then there will always be a minority of cells with parameters that locate them near the boundary with the more favorable phenotype so that even mutations with small-size effects can result in movement into a qualitatively different phenotypic region that is favorable. Thus, evolvability coexists with robustness. A statistical approach within the Design Space framework could be used to test these speculations.

The results in **Figure 6**, which represent the most extreme bottleneck with a single founder cell, suggest that most of the phenotypes are regenerated with sizable cell numbers within the initial growth phase. A stochastic approach could be used to study the long-term fate of the remaining phenotypes whose population sizes are < ∼ 100 cells, each of which is retained or lost in each generation.

In this paper, the phenotype-centric approach provides a novel theoretical framework to pose and answer questions of phenotype-specific mutation rates and ranking of phenotype frequencies in the population under non-selecting and selecting conditions. This framework makes a key distinction between ‘*entropy increasing/entropy decreasing’* mutations, which cause genetically-determined parameter values to change in the direction of an increase/decrease in entropy, and ‘*beneficial/detrimental’* mutations, which cause the integrated activities of the entire system to change in the direction of an increases/decreases in phenotype fitness. The two causes are separable. The importance of the distinction can be exemplified by considering the consequence for a population evolving in a temperature gradient (Zhang et al., 2018; Wooliver et al., 2020).

In an idealized case, if the population finds itself in a new environment with a higher temperature than the one in which it was previously adapted, the binding of a regulator will now be less effective (higher temperature implies looser binding). The fitness of the organisms will typically decrease. An *entropy-decreasing* mutation causing tighter binding of the regulator can cause an improvement in fitness. Conversely, an *entropy-increasing* mutation causing an even looser binding can cause a further reduction in fitness. The argument is different if the population finds itself in a new environment that has a lower temperature. The binding of the regulator will now be too tight (lower temperature implies stronger binding). The fitness of the organisms will typically decrease. Now, an *entropy-increasing* mutation that causes a looser binding of the regulator can improve fitness. Conversely, an *entropy-decreasing* mutation that causes an even tighter binding can cause a further reduction in fitness. Thus, depending on the environmental condition, an entropically probable mutation at the level of the molecular mechanism can cause either a beneficial or detrimental effect on fitness at the level of an integrated system (**Figures 3 & 7**). These same distinctions provide a mechanistic context for interpreting the large differences in frequency of positive-effect mutations that have been discussed by Bondel, et al. (2019).

The theoretical framework based on Design Space analysis can distinguish and quantify various phenomena. For example, it distinguishes among three contributions to phenotype-specific mutation rates: phenotype robustness (volume in Design Space), size effect and directional bias, and selection (**Figure 3**); distinguishes among three contributions to the equilibrium distribution with neutral fitness effects(**Eqn. 9**): mutation alone and selection alone, which nearly balance, and mutation-x-selection (mutations generated specifically by the selected phenotype), which is only significant with extremely strong selection (**Figure 4**); quantifies the different time scales of evolution between equilibria under selecting and non-selecting conditions (**Figure 5**).

### Summary of Results Old and New

The findings in **RESULTS** agree with many well-known phenomena in theoretical population genetics. Examples include stronger selection is needed to counteract higher mutation rates, evolution can be faster with higher mutation rates, positive-effect mutations are rare in well adapted systems and small-effect mutations are common, and the characteristic distributions observed in *directional* (Darwin, 1859; Mitchell-Olds et al., 2007) and *stabilizing* (Charlesworth et al., 1982; Campbell & Reece, 2002) selection; *mutation-selection balance* (Barton, 2007; Lynch, 2010), and cryptic variation under non-selecting conditions (Paaby et al., 2014; Zheng et al., 2019).

However, in all these cases the Design Space framework provides a more nuanced understanding of their underlying molecular mechanisms with phenotype-specific mutation playing a role in each. For example, the phenotype distribution with no size effect, directional bias or differences in growth rate under the non-selecting condition, which might be expected to produce a uniform distribution of mutant effects, is weakly directional even though no selection is involved (**Figure 3A**, Blue); the causal fitness characteristic is the robustness (polytope volume) of phenotypes with phenotype #6 dominating. The phenotype distribution with size effect and directional bias but no differences in growth rate under the non-selecting condition is more strongly directional even though no selection is involved (**Figure 3A**, Black), with phenotype #6 dominating; the causal fitness characteristics are robustness and entropy. Although the phenotype distribution with size effect, directional bias and protein burden differences in growth rate under the non-selecting condition may also appear to be directional (**Figure 3B**, Red), with phenotype #11 dominating, it is actually balancing since the causal fitness characteristics are a balance between protein burden differences in growth rate in one direction and entropy in the other. Furthermore, the point of balance is a function of the general mutation rate *m*, which is 10^−7^ in this case. With a higher general mutation rate *m*, the balance shifts in favor of entropy (**Figure 3B)**, and as *m* approaches 10^−4^, entropy dominates to such an extent that the distribution suggests directional selection. The phenotype distribution with size effect, directional bias and protein burden differences in growth rate under the selecting condition is a more complex balancing selection (**Figure 3C**, Black), with phenotype #7 dominating; the causal fitness characteristics are a balance between protein burden differences in growth rate in one direction and the selective advantage of oscillation and entropy in the other. The general mutation rate (*m* = 10^−7^ in this case) also plays a causal role in the balance. The distribution of cryptic variation present under non-selecting conditions differs, depending on whether the fitness effects of mutations are neutral (**Figure 3A**,Black) vs. near-neutral (**Figure 3B**, Red). Although it is difficult to measure such small differences in fitness experimentally, the resulting distributions are markedly different, as are the results under selection (**Figure 7**). The causal fitness characteristics involved in the balance are entropy, protein burden differences in growth rate, and genomic mutation rate; the first is dominant in the neutral case, the second is dominant in the near-neutral case, and the third is capable of eliminating the distinction between neutral and near-neutral at sufficiently high rates (**Figure 3B**, Blue).

Other results are new, e.g., there is an optimal mutation rate for each phenotype (**Figures 3 & S7**); the percentage of positive effect mutations is smaller when equilibrium is dominated by phenotypes with high entropy and larger when dominated by those with low entropy (**Figure 7**); evolution is slower in the former an faster in the latter; there are many changes in population rank with weak selection (**Figure 5**), and few with strong selection (not shown); a non-selected phenotype can increase (without hitch-hiking) as an indirect result of selection for a different phenotype connected by a high phenotype-specific mutation rate (**Figure S8**); and back calculation of selection coefficients is possible from well characterized distributions (**Figure 7**). We also provide evidence suggesting that experimental evolution in chemostats can be used to experimentally test predictions made possible by the phenotype-centric theory (**Figure 6**).

We return to the fundamental question raised at the outset and ask, what is the relevant distribution of phenotype frequencies to consider from which there is evolution of new phenotypes? This is still an open question. New phenotypes will grow to dominance when the population suddenly finds itself in a selecting condition because of a change in genotype or a change in environment. The results for the clock model suggest that the equilibrium distribution of the full repertoire in the non-selecting condition with neutral fitness effects, might be most relevant to consider (**Figure 7**). However, even small differences from neutrality that are experimentally undetectable, such as protein burden effects, can result in a marked difference in the distribution (**Figure 3**) that argue against its relevance in the case of the natural *lac* operon.

Finally, it should be noted that although we have emphasized qualitatively-distinct phenotypes, *quantitative* variants exist within each phenotypic region in System Design Space. Thus, the phenotype-centric approach also provides the opportunity to explore finer changes in quantitative characteristics such as frequency, phase and amplidude of the oscillations within the region of phenotype #7 (Lomnitz & Savageau, 2013). Such results would be relevant to the work of Ouyang et al. (1998) showning that mutants with small changes in frequency of the cyanobacteria circadian clock experience negative selection when their frequency differs from that of the environmental light-dark cycle.

## MATERIALS AND METHODS

Methods developed in this work are described in the section DERIVATION OF PHENOTYPE-SPECIFIC MUTATION RATE CONSTANTS AND POPULATION DYNAMIC EQUATIONS. Associated computational tools with further details can be accessed through the Design Space Toolbox v.3.0, which is freely available for all major operating systems via Docker. After Docker has been installed on your system, running the following commands on a terminal window will provide access to the software:

1. docker pull savageau/dst3
2. docker run -d -p 8888:8888 savageau/dst3
3. Access the software by opening the address http://localhost:8888/ on any internet browser.

Please refer to Valderrama-Gómez et al. (2020) for detailed installation instructions and troubleshooting. Several IPython notebooks are provided to reproduce figures in the main text and supplementary information. These notebooks can be found within the Docker image (savageau/dst3) under the directory **/Supporting_Notebooks/MBE**.

## ACKNOWLEDGMENTS

This work was supported in part by a grant from the US National Science Foundation Grant number MCB 1716833.

