## Supplemental Information for "Molecular Systems Predict Equilibrium Distributions of Phenotype Diversity Available for Selection"

Miguel A. Valderrama-Gómez[1] and Michael A. Savageau[1,2,*]

[1]Department of Microbiology & Molecular Genetics, and [2]Department of Biomedical Engineering, University of California, Davis, One Shields Avenue, Davis, CA 95616 USA

Keywords: biochemical systems theory / system design space / phenotype-centric modeling / theoretical population genetics / Fisher's Geometric Model / evolutionary dynamics / circadian clock circuitry

### TABLE OF CONTENTS


| SECTIONS | TITLE | PAGE |
|---|---|---|
| ***S1*** | ***Brief Summary of Design Space Concepts*** | 3 |
| *S1.1* | *Biochemical Phenotype* | 3 |
| *S1.2* | *Phenotypic Repertoire* | 4 |
| *S1.3* | *Phenotypic Properties* | 5 |
| ***S2*** | ***Biochemical Kinetic Equations*** | 5 |
| ***S3*** | ***Generalized Mass Action System of Equations*** | 6 |
| ***S4*** | ***Methods for Calculating Phenotype Volumes*** | 7 |
| ***S5*** | ***Rescaling the Molecular Model Reveals an Invariant System Design Space*** | 7 |
| ***S6*** | ***Comparison of Methods for Calculating Phenotype Volumes*** | 9 |
| ***S7*** | ***Geometry of Tracks, Volumes and Distance in System Design Space*** | 10 |
| ***S8*** | ***Distance Between Centers of Volume*** | 11 |
| ***S9*** | ***Directional Bias Parameter*** | 12 |
| ***S10*** | ***Systemic Effects of Mutation and Design Principles*** | 13 |
| ***S11*** | ***Comparisons to Experimental Studies*** | 14 |
| ***S12*** | ***Temporal Response at Higher Mutation Rates*** | 17 |
| FIGURES & TABLES | | |
| **Figure S1** | **Importance of orientation for phenotype volume in System Design Space** | 7 |
| **Figure S2** | **Design space analysis of the dimensionless version of the mechanistic model** | 9 |
| **Figure S3** | **Comparison of four approximate methods for calculating phenotype volumes in system design space** | 10 |
| **Figure S4** | **The parameters defining biochemical phenotypes of a molecular mechanism are located within distinct convex polytopes separated by hyperplanes in System Design Space** | 11 |
| **Figure S5** | **Comparison of the point-by-point and centers of volume methods for calculating phenotype-specific mutation rates** | 12 |
| **Table S1** | **The influence of distance between phenotypes and bias in the direction of parameter change on the probability of mutation** | 13 |
| **Figure S6** | **Steady-state induction characteristic for the model characterized in Figure 2A** | 15 |
| **Figure S7** | **Distribution of qualitatively-distinct phenotypes for constructed mutants distinct** | 16 |
| **Figure S8** | **Temporal response in relative frequency of phenotypes following imposition of the selecting condition** | 18 |
| REFERENCES | | 18 |

### SUPPLEMENTAL INFORMATION

#### ***S1. Brief Summary of Design Space Concepts***

The System Design Space approach enables a novel 'phenotype-centric' modeling strategy that is radically different from the conventional 'simulation-centric' approach (Valderrama-Gómez et al., 2018). It enumerates the phenotypic repertoire algebraically, without a priori knowledge of the kinetic parameter values, and then predicts parameter values for phenotypes of interest. This is the inverse of the simulation-centric approach in which parameter values must first be determined (by measurement, estimation, sampling etc.), and then the phenotypic repertoire can be sampled by computer simulation. For a detailed description of this material, the interested reader is referred to Valderrama-Gómez et al. (2020), which also describes the methods for automated analysis based on biochemical phenotypes. These concepts also will be illustrated in the context of the specific case study in the main text. Surprisingly, as noted in the main text, much of what can be learned about the system depends largely on the architecture of the model. By model architecture we mean molecules, interactions among them, and the signs of the interactions.

##### *S1.1. Biochemical Phenotype*

The scope of Biochemical Systems Theory includes mechanistic models governed by fundamental rate laws. These are the power functions of chemical kinetics and the rational functions of biochemical kinetics. These functions are integrated into a network based on Kirchhoff's node law. The result is a system of differential-algebraic equations that can be transformed, without loss of generality, into a corresponding generalized mass action system (GMA-system) of equations (Savageau & Voit, 1987). A sub-system (S-system) of differential algebraic equations is defined by a dominant positive and negative term from each of the GMA-system equations. Those S-systems possessing a self-consistent steady state solution and passing the test for dominance in the full GMA-system, provide the following definitions of phenotype based on Biochemical Systems Theory. A *phenotype* is the set, or sets, of concentrations and fluxes corresponding to a valid combination of dominant processes functioning within an intact system. A *qualitatively distinct phenotype* is the characteristic phenotype that exists throughout a region of validity (polytope) in parameter space. A phenotypic repertoire is the collection of qualitatively distinct phenotypes integrated into a space-filling structure in parameter space.

This definition of a biochemical phenotype involves all the variables and parameters of the full system. It also provides rigorously defined boundaries in logarithmic parameter space delineating a high-dimensional volume (or polytope), characterized by linear hyperplanes and vertices determined by their intersections, within which the parameter values yield the same *qualitatively distinct biochemical phenotype*. The inequalities defined by these boundaries provide specific design principles for the realization of specific phenotypes with specific behaviors. Thus, biochemical phenotypes are characterized at several interrelated levels by *mechanisms* (those portions of the system's mechanisms that are being exercised under specific genetic and environmental conditions), *equations* (a specific S-system equation), *geometry* (volumes and boundaries in parameter space), *design* (specific design principles), and *behavior* (both qualitative and quantitative).

It must be emphasized that 'dominance' in the context of System Design Space means simply that one of many nodal fluxes, mechanistic contributions, or terms in the dynamic balance equations is mathematically larger than the others (by however small an amount); it does not imply that the others are being omitted, neglected, or ignored. In a sense, all of them are being determined to verify that the assumption of dominance is in fact true. Thus, all the system's parameters and dynamic variables are involved in determination of the S-system equation and the boundaries for its validity. It may be counterintuitive, but all the system's concentrations and fluxes are predicted in this form of system deconstruction. The concentrations are predicted directly by the solution of the S-system and the fluxes, dominant and non-dominant, are predicted by a simple secondary calculation from the concentrations.

*S1.2. Phenotypic Repertoire*

Qualitatively distinct phenotypes, as represented by their polytope volumes, fill the entire space of parameter values, decomposing this space into a finite number of discrete 'chunks'. The enumeration of the full repertoire of qualitatively distinct phenotypes is performed automatically by DST3. The phenotypes in the repertoire are assigned a specific number and signature, which identifies the corresponding dominant terms in each of the original equations.

*S1.3. Phenotypic Properties*

DST3 currently predicts a large number of phenotype characteristics automatically, including (1) a set of *nominal parameter values* for the realization of each phenotype, (2) the *global tolerances* to change of each parameter before there is a qualitative change in phenotype, the product of which provides an under-estimate of the phenotype's polytope volume, (3) the *eigenvalues* characterizing the dynamic behavior of the phenotype, (4) *signal amplification (logarithmic gain)* factors between specific genetic and environmental input signals and the system's concentrations or fluxes as output signals, (5) the *volume* of parameter space enclosed by each phenotypic polytope, (6) the *centroid* of each phenotypic polytope and, as we will now show, (7) *phenotype-specific mutation rate constants*.

***S2. Biochemical Kinetic Equations***

The equations used to represent the precursor clock module are based on the foundation of fundamental biochemical kinetics, which has broad general applicability as indicated by the vast majority of biochemical models that are of this type (Chelliah et al., 2013). The 4-stage model in **Figure 1** is a simplification of models that have been used to represent biochemical oscillators (Stricker et al., 2008; Lomnitz and Savageau, 2014) that include an additional stage for protein folding or maturation. Although this makes for a 6-stage model, two of the stages can be considered in quasi-steady state and effectively combined with others because the two represent fast processes (mRNA degradation), leaving 4-stages that are the slower temporally-dominant processes (protein degradation, protein folding). Thus, combining transcription and translation yields the same 4-stage model in **Figure 1**, but with the original variables *mP* (mRNA encoding the positive transcription factor) and *mN* (mRNA encoding the negative transcription factor) now representing the concentration of unfolded P and N protein.

$$\frac{dmP}{dt} = \frac{\alpha_{mP\max} + \alpha_{mP\min}\left(\frac{N}{K_N}\right)^n}{1+\left(\frac{N}{K_N}\right)^n} - \beta_{mP} mP \qquad \alpha_{mP\max} > \alpha_{mP\min} \tag{S1}$$

$$\frac{dP}{dt} = \alpha_P mP - \beta_P P \tag{S2}$$

$$\frac{dmN}{dt} = \frac{\alpha_{mN\min} + \alpha_{mN\max}\left(\frac{P}{K_P}\right)^p}{1+\left(\frac{P}{K_P}\right)^p} - \beta_{mN} mN \qquad \alpha_{mN\max} > \alpha_{mN\min} \tag{S3}$$

$$\frac{dN}{dt} = \alpha_N mN - \beta_N N \tag{S4}$$

The $K_i$ parameters reflect the binding of the regulators to their promoter targets; the kinetic orders $n$ and $p$ reflect the number of binding events (cooperativity) in the interactions; the $\alpha_{i\max}$ and $\alpha_{i\min}$ parameters are the maximum and minimum rates of mRNA/protein expression; the $\alpha_i$ parameters are the first-orter rate constants for protein folding, and the $\beta_i$ parameters are first-order rate constants for loss.

***S3. Generalized Mass Action System of Equations***

These equations, recast as an equivalent GMA-system (Savageau & Voit, 1987), can be expressed as a set of differential-algebraic equations in the syntax of the Design Space Toolbox (Lomnitz and Savageau, 2016).

$$mP. = amP\max*(DP\wedge -1) + amP\min*(KN\wedge -n)*(N\wedge n)*(DP\wedge -1) - bmP*mP \tag{S5}$$

$$P. = aP*mP - bP*P \tag{S6}$$

$$mN. = amN\min*(DN\wedge -1) + amN\max*(KP\wedge -p)*(P\wedge p)*(DN\wedge -1) - bmN*mN \tag{S7}$$

$$N. = aN*mN - bN*N \tag{S8}$$

$$0 = 1 + (KN\wedge -n)*(N\wedge n) - DP \tag{S9}$$

$$0 = 1 + (KP \wedge -p) * (P \wedge p) - DN \qquad \textbf{(S10)}$$

with parametric constraints

$$amPmax > amPmin \qquad \textbf{(S11)}$$

$$amNmax > amNmin \qquad \textbf{(S12)}$$

##### *S4. Methods for Calculating Phenotype Volumes*

Accurate volumes of the phenotypes in Design Space can be calculated by means of vertex enumeration methods (Avis 2000, Barber et al. 1996), which work well for small systems but currently do not scale well for large systems. There are other readily available means to automatically estimate approximate phenotype volumes such as those in **Figure S3** that do scale well.

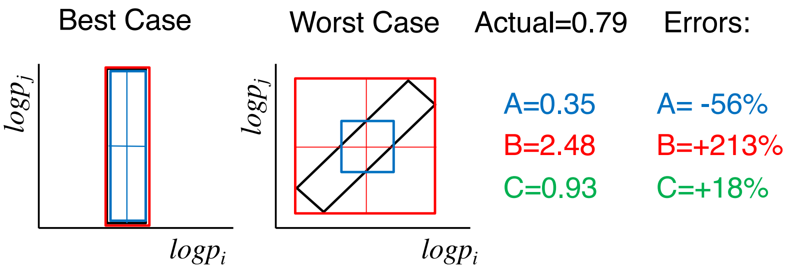


**Figure S1. Importance of orientation for phenotype volumes in System Design Space.** Illustration involving a narrow rectangle. (**Left Panel**) Best case in which the sides of the rectangle are parallel to the axes. The volume estimated by the product of parameter tolerances (Blue) and by the bounding box (Red) methods are exact and identical. (**Right Panel**) Worst case in which the sides of the rectangle are rotated between the axes. (**A**) The volume estimated by the product of parameter tolerances method (Blue) is an under-estimate. (**B**) The volume estimated by the bounding box method (Red) is an over-estimate. (**C**) The volume estimated by the geometric mean of the two methods (Green) provides a better approximation to the true volume.

##### *S5. Rescaling the Molecular Model Reveals an Invariant System Design Space*

Dimensionless versions of the system variables can be defined as $m\pi = mP/[\alpha_{mP\max}/\beta_{mP}]$, $\pi = P/[K_P]$, $m\nu = mN/[\alpha_{mN\max}/\beta_{mN}]$ and $\nu = N/[K_N]$, and dimensioless versions of the system parameters can be defined as $\rho_P = \alpha_{mP\max} / \alpha_{mP\min} > 1$, $\rho_N = \alpha_{mN\max} / \alpha_{mN\min} > 1$, $m\pi_{\max} = 1$, $m\nu_{\max} = 1$ and

$$\pi_{\max} = \frac{1}{K_P}\left(\frac{\alpha_{mP\max}}{\beta_{mP}}\right)\left(\frac{\alpha_P}{\beta_P}\right) \quad \textbf{(S13)}$$

$$\nu_{\max} = \frac{1}{K_N}\left(\frac{\alpha_{mN\max}}{\beta_{mN}}\right)\left(\frac{\alpha_N}{\beta_N}\right) \quad \textbf{(S14)}$$

The dimensionless version of the 12-parameter model (**Eqns. S1-S4**) can then be written as

$$\frac{1}{\beta_{mP}}\frac{dm\pi}{dt} = \frac{1+\frac{1}{\rho_P}(\nu)^n}{1+(\nu)^n} - m\pi \quad \textbf{(S15)}$$

$$\frac{1}{\beta_P}\frac{d\pi}{dt} = \pi_{\max} m\pi - \pi \quad \textbf{(S16)}$$

$$\frac{1}{\beta_{mN}}\frac{dm\nu}{dt} = \frac{\frac{1}{\rho_N}+(\pi)^p}{1+(\pi)^p} - m\nu \quad \textbf{(S17)}$$

$$\frac{1}{\beta_N}\frac{d\nu}{dt} = \nu_{\max} m\nu - \nu \quad \textbf{(S18)}$$

The 4-parameter dimensionless model (**Eqns. S15-S18**) is goverened by two parameters, $\rho_P$ *and* $\rho_N$ that shape the phenotype polytopes and two, $\pi_{\max}$ *and* $\nu_{\max}$ that only translate the position of the nominal operating point within an invariant System Design Space. As a consequence, 10 of the original 12 parameters have only a proportional effect on translating the invariant System Design Space.

Note that the axes in **Figure S2A** have a negative sign because the values for dimensional groups $\nu_{\max}$ and $\pi_{\max}$ (**Eqns. S13** and **S14**) are inversely related to the values for $K_N$ and $K_P$ in **Figure 2A**. The damped oscillations in **Figure S2B** are identical to those in **Figure 2C**, except for differences in scale for the variables. Finally, time in **Figure S2B** also can be scaled by a factor of 1/3 to produce an oscillation with a 24-hour period (**Figure 2C**).

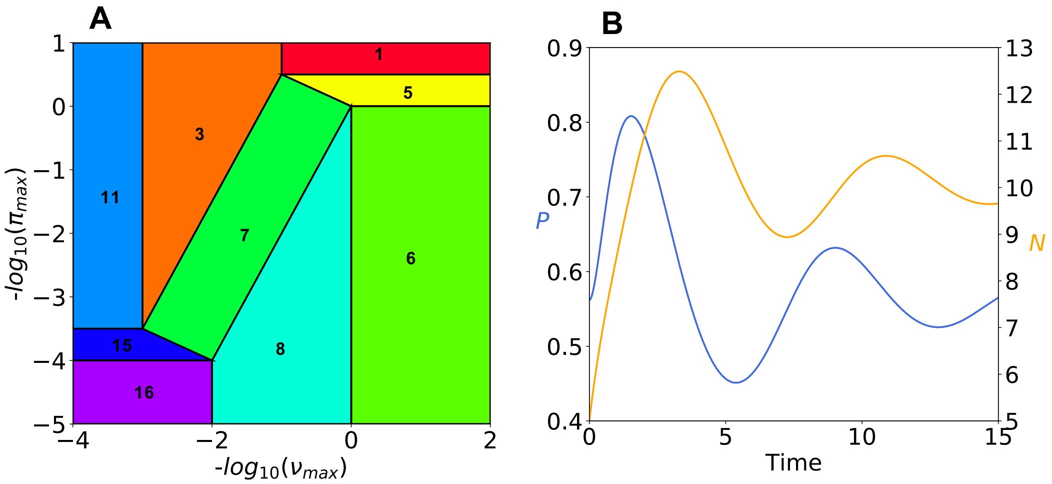


**Figure S2. Design space analysis of the dimensionless version of the mechanistic model. (A)** Enumerated qualitatively-distinct phenotypes identified by color for the invariant System Design Space. **(B)** Dynamic behavior of the oscillatory phenotype #7. Concentrations of P and N are plotted against time. Initial conditions are: mP = 0.0100018352905; P = 0.561900859018; mN = 0.315970420699; N = 5.0. Figures generated with the following parameter values: aNmax = 0.0316; aPmax = 0.0178; rN = 10.0; rP = 10000.0; Kinetic order(s): n=2, p=2; Parametric constraints: rP >1, rN >1.

##### *S6. Comparison of Methods for Calculating Phenotype Volumes*

Of the three methods described in **Figure S1**, the one determined by the product of the global tolerances from the central value for each parameter gives an underestimate, another determined by the product of the largest extents for each of the parameters (the so-called bounding box) gives an overestimate, and a third determined by the geometric mean of the first two approximate estimates can be used as an improved estimate. The differences among the three are illustrated graphically in **Figure S1**. A fourth method under development involves the product of the track widths and the tolerance of the mutated parameter along the track midway between the values of the orthogonal parameters (red line in **Figure S4B,C,D**). The results of comparisons among the four methods are summarized in **Figure S3**

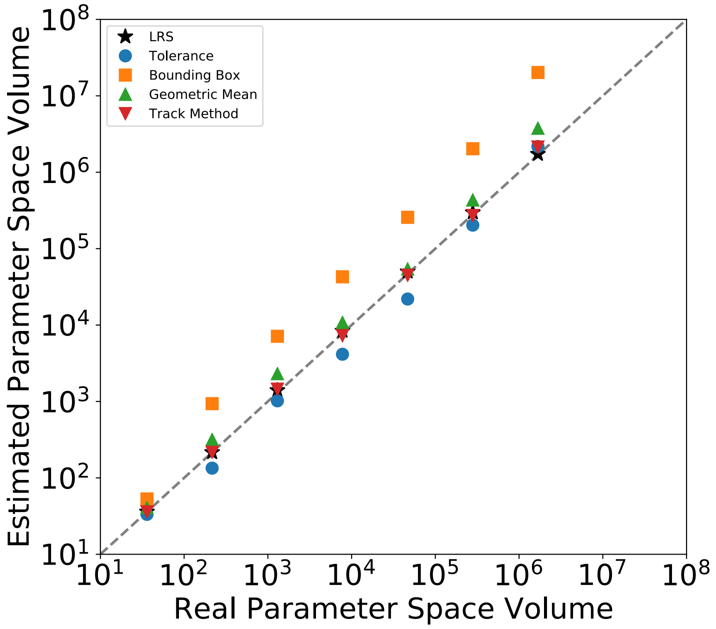


**Figure S3. Comparison of four approximate methods for calculating phenotype volumes in System Design Space. The standard is the vertex LRS method.** The average absolute errors are LRS (5%), track (8%), tolerance (32%), geometric mean (53%) and bounding box (494%).

#### *S7. Geometry of Tracks, Volumes and Distance in System Design Space*

A two-dimensional illustration of the more general situation is shown in **Figure S4A**, where the areas correspond to the *n*-dimensional polytope volumes of the phenotypes. This illustration involves two parameters on the x- and y- axis, three phenotypes depicted in three colors, seven vertices with four on the corners representing the universe of parameter values and three on the edges, six linear hyper-planes with four bounding the universe of parameter values and two intersecting the parameter space, four tracks with two for each of the parameters, and five volumes with one phenotype having a single volume and two phenotypes having two sub-volumes each.

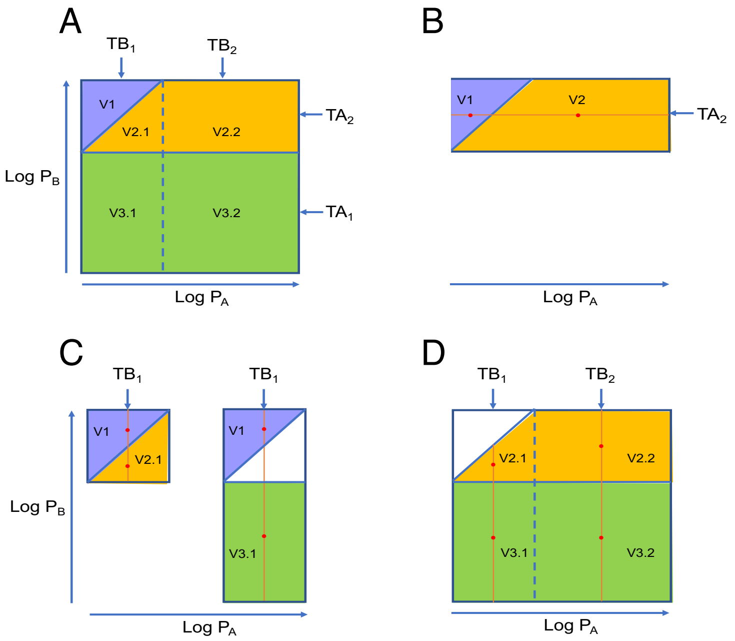


**Figure S4. The parameters defining biochemical phenotypes of a molecular mechanism are located within distinct convex polytopes separated by hyperplanes in System Design Space. (A)** In this two-dimensional illustration there are three phenotypes (#1, #2 and #3) with different polytope volumes (areas): V1, V2=V2.1+V2.2 and V3=V3.1+V3.2. **(B)** Mutations between phenotypes #1 and #2 due to changes in parameter $P_A$ occur along *mutation track* TA2. **(C, Left)** Mutations between phenotypes #1 and #2 due to changes in parameter $P_B$ occur along mutation track TB1. **(C, Right)** Mutations between phenotypes #1 and #3 due to changes in parameter $P_B$ occur along mutation track TB1. **(D)** Mutations between phenotypes #2 and #3 due to changes in parameter $P_B$ occur along mutation tracks TB1 and TB2. The *track widths* are defined by the difference between the least upper and the greatest lower *bounding box coordinates* for the *orthogonal parameter* of the donor and recipient phenotype regions. The *centroid* in each region is given by ½ the track width (red line) and ½ the *parameter tolerance* along the red line (red dot). Phenotype-specific mutation rates between donor and recipient phenotypes are determined by the differences between their centroids and the volume of the recipient phenotype.

##### *S8. Distance Between Centers of Volume*

The average mutation rate can be determined by dense sampling of the individual point-by-point differences between phenotypes and averaging. Alternatively, the same result can be obtained using only the differences between the centers of volumes of the phenotypic regions (**Figure 2**).

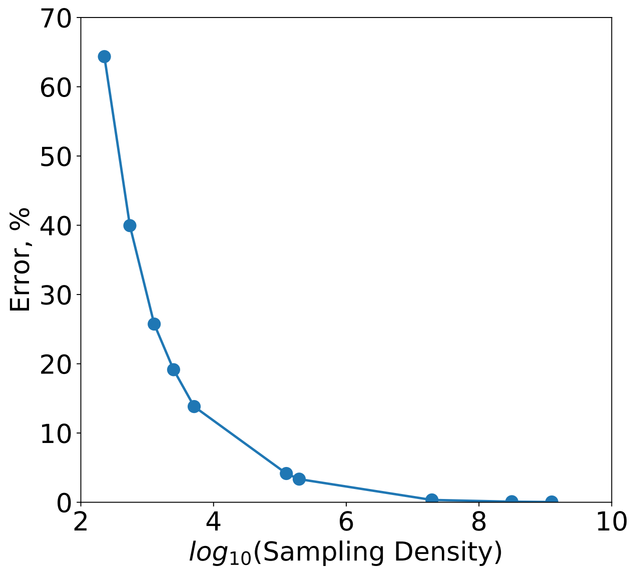


**Figure S5. Comparison of the point-by-point and centers of volume methods for calculating phenotype-specific mutation rates.**

The point-by-point method produces an estimate that converges asymptotically to the centers of volumes method as the density of sampling increases (**Figure S5**).

***S9. Directional Bias Parameter***

Entropy dictates that parameter changes are more probable in one direction vs. the alternative. The probability of change in the rate constant for mRNA/protein degradation is greater in the direction of increase (decrease in stability or lifetime) and lesser in the direction of decrease (increase in stability or lifetime). Similarly, the probability of change in the rate constant representing the minimum rate of transcription/translation is greater in the direction of increase (leaky promoter) and lesser in the direction of decrease (tight promoter). On the other hand, the probability of change in the rate constant representing the maximum rate of transcription/ translation is greater in the direction of decrease (weak promoter) and lesser in the direction of increase (strong promoter). The probability of change in a dissociation equilibrium constant is greater in the direction of increase (loose binding) and lesser in the direction of decrease (tight binding). These types of directional bias are represented mathematically in **Table S1**.

**Table S1. The influence of distance between phenotypes and bias in the direction of parameter change on the probability of mutation.**

| Parameter | Direction of Parameter Change | |
|---|---|---|
| | Increase | Decrease |
| $K$ | $\exp[-s/(\lambda*\delta)]$ | $\exp[-s/(\lambda/\delta)]$ |
| $\beta$ | $\exp[-s/(\lambda*\delta)]$ | $\exp[-s/(\lambda/\delta)]$ |
| $\alpha$ | $\exp[-s/(\lambda/\delta)]$ | $\exp[-s/(\lambda*\delta)]$ |
| $\alpha_{max}$ | $\exp[-s/(\lambda/\delta)]$ | $\exp[-s/(\lambda*\delta)]$ |
| $\alpha_{min}$ | $\exp[-s/(\lambda*\delta)]$ | $\exp[-s/(\lambda/\delta)]$ |

Parameters: equilibrium dissociation constants, $K$, first-order rate constants for the degradation of mRNA or protein, $\beta$, first-order rate constants for translation, $\alpha$, and max/min values for rates of transcription, $\alpha_{\max}/\alpha_{\min}$. The size, $s$, is the magnitude of the difference between values of the given parameter. The size scale, $\lambda$, and directional bias, $\delta$, are the two parameters of the distribution.

##### *S10. Systemic Effects of Mutation and Design Principles*

In **RESULTS**, we provide an example of a simple *system design principle* involving only changes in the two equilibrium dissociation constants. However, changes such as these also will have a systemic influence on the expression of both activator and repressor. When the interactions among all the parameters and variables are considered, the boundaries between phenotypes are determined solely by the parameter values (results of DST3 *Analyze Case* command for Case #7, results not shown). The result is a subtle *system design principle* defined by the four boundaries

$$1 < \left[\frac{1}{K_P}\left(\frac{\alpha_{mP\max}}{\beta_{mP}}\right)\left(\frac{\alpha_P}{\beta_P}\right)\right]^{-\frac{np}{1+np}} \left[\frac{1}{K_N}\left(\frac{\alpha_{mN\max}}{\beta_{mN}}\right)\left(\frac{\alpha_N}{\beta_N}\right)\right]^{-\frac{n}{1+np}} < \rho_P \qquad \textbf{(S19)}$$

$$1 < \left[\frac{1}{K_P}\left(\frac{\alpha_{mP\max}}{\beta_{mP}}\right)\left(\frac{\alpha_P}{\beta_P}\right)\right]^{\frac{p}{1+np}} \left[\frac{1}{K_N}\left(\frac{\alpha_{mN\max}}{\beta_{mN}}\right)\left(\frac{\alpha_N}{\beta_N}\right)\right]^{-\frac{np}{1+np}} < \rho_N \qquad \textbf{(S20)}$$

or in terms of the dimensionless parameter groups defined in **Section S5**

$$1 < \left[\pi_{\max}\right]^{-\frac{np}{1+np}} \left[\nu_{\max}\right]^{-\frac{n}{1+np}} < \rho_P \qquad \textbf{(S21)}$$

$$1 < \left[\pi_{\max}\right]^{\frac{p}{1+np}} \left[\nu_{\max}\right]^{-\frac{np}{1+np}} < \rho_N \qquad \textbf{(S22)}$$

Violation of one of these four inequalities results in a mutation from phenotype #7 to one of its adjacent neighbors. The specific example of a transition from phenotype #7 to #5 discussed in **RESULTS** corresponds to a violation of the left inequality in **Eqn.** (**S19 or S21**). The most effective mutations promoting this transition are among the parameters of $\pi_{\max}$ since they are raised to a higher power ($p = 2$), e.g. if a 100-fold change in $\nu_{\max}$ were necessary then only a 10-fold change in $\pi_{\max}$ would suffice.

#### *S11. Comparisons with Experimental Studies*

There have yet to be any specific experimental studies designed to test the detailed predictions this theory generates. However, there are existing reports in the literature that could be considered, with the appropriate caveats. There are studies that attempt to infer the distribution of mutation effects at the level of specific functions (e.g., Markiewicz et al., 1994) or at the level of the whole genome (e.g., Bondel et al., 2019). We address just these two examples here.

One of the best studied systems, for which the fitness characteristics of a specific function have been examined, is the LAC repressor of *E. coli* binding to its specific recognition site in the DNA (Markiewicz et al., 1994). Both the LAC repressor in the inducible system and the N repressor in the oscillatory system, have three qualitatively distinct phenotypes: a 'wild-type' phenotype capable of performing the necessary function (induction or oscillation), an entropically more probable phenotype consisting of 'negative-effect' constitutive mutants, and an entropically less probable phenotype consisting of 'positive-effect' super-repressed mutants (**Figure S6**). In both wild-types, the binding of repressor must occupy a "Goldilocks" state that is just right, not too strong and not too weak, and mutations in either direction away from this state are dysfunctional.

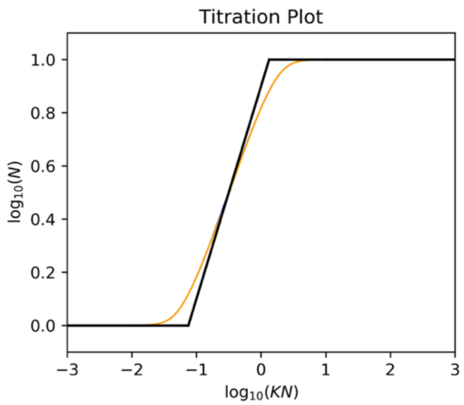


**Figure S6. Steady-state induction characteristic for the model characterized in Figure 2A.** The system exhibits three phenotypic regions in response to mutational change in the equilibrium dissociation constant $K_N$. Changes in the N gene expression are super-repressed ($K_N < 0.1$, phenotypes #3 and #11), regulated oscillatory ($0.1 < K_N < 1.1$, phenotype #7), and constitutive ($K_N > 1.1$, phenotypes #6 and #8). S-system approximate values (Black) and full system values (Gold). The calculations shown correspond to the system defined by **Eqs. S5** to **S12** which is parameterized using values from **Table 2** for all parameters except for $K_N$.

Markiewicz et al. (1994) did a fine-structure map of mutations in the *lacI* gene by using alternative codon assignments with suppressors to change amino acid residues at each of 328 positions in the LAC repressor protein. Since the repressor function in our model system does not involve the binding of an inducer, for purposes of comparison, we will treat positions in the LAC protein that involve inducer binding as being tolerant to substitutions, as variations are tolerated in related proteins (Markiewicz et al, 1994). Their data show that mutations at ~ 67% of the positions had a 'zero effect' (tolerant to substitutions), 31% had a 'negative effect' (decrease in DNA binding), and ~2% had a 'positive effect' (increase in DNA binding).

If we select values of $\lambda = 0.6$ and $\delta = 1.85$, we can calculate the corresponding values for the probability of mutations in the N gene remaining within its wild-type oscillatory phenotype ('zero effect': #7), leaving for phenotypes with decreased DNA binding ('negative effect': #6, #8), and leaving for phenotypes with increased DNA binding ('positive effect': #3, #11). The resulting values for the distribution are then ~ 67% zero-effect, ~31% negative-effect, and ~2% positive-effect, which matches the experimental results.

The degree of selection required to maintain this phenotype distribution against the mixed fitness background with *m* = 1.0E-07 as shown in **Figure 7B** is 4.2E-05 %. If this selection is removed, the distribution evolves toward the equilibrium condition shown in **Figure 7B** (at $\mu_7^* = 1$), **Figure 5B** and **Figure S7** (at *m* = 1.0E-07). Even at the optimum mutation rate in the non-selecting condition for phenotype #7, *m* = 2.2E-06, the equilibrium distribution with mixed fitness effects is very different from that with neutral fitness effects as shown in **Figure 7A** (at $\mu_7^* = 1$), **Figure 5A** and **Figure S7** (at *m* = 1.0E-04). The equilibrium phenotype distribution in either case is very different from that of wild-type LAC repressor selected in nature (**Figure S7**, flat lines.

In contrast to the focus on fitness characteristics of a specific function, Bondel et al. (2019) used a genome-scale approach involving a mutation-accumulation experiment, with no

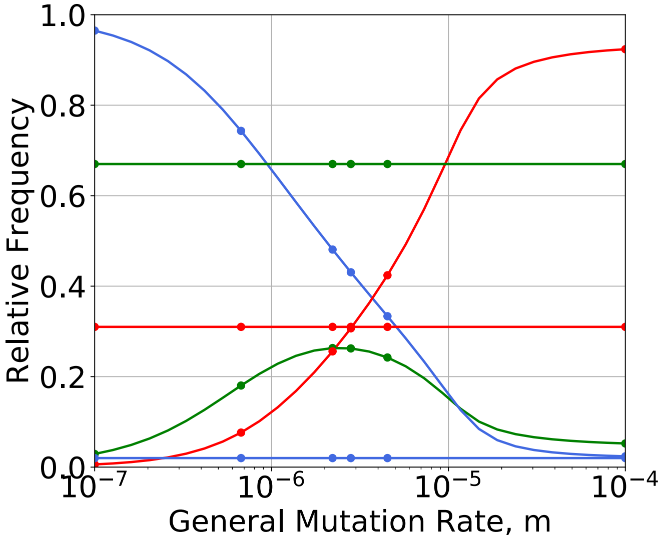


**Figure S7. Distribution of qualitatively-distinct phenotypes for constructed mutants**. Phenotype distributions under non-selecting conditions at equilibrium as a function of the general mutation rate *m* (curved lines) and at non-equilibrium (time-zero measurements from the system having previously been under selecting conditions) independent of *m* (flat lines). Phenotypes: constitutive (Red), oscillatory (Green), and super repressed (Blue). The distribution for the oscillatory phenotype is optimal at *m* = 2.2E-6. Dots, from left to right, represent optimal mutation rates for phenotypes #11, #3, #7, #15, #16, #8, #6, #5 and #1. Note that phenotypes #16 and #8 share the same optimum. This is also true for phenotypes #6, #5 and #1.

attempt to identify the underlying functional mechanisms, to estimate the distribution of fitness effects based on the growth rate of *Chlamydomonas reinhardtii*. In a three-category statistical model, they estimated that ~ 88% of the mutations were zero-effect, ~7% were negative-effect, and ~5% were positive-effect. In their double-sided gamma distribution model, they found negative-effect mutations six-fold more prevalent than positive-effect mutations for mutations with absolute fitness > 1%; the percentages in this case are then probably closer to 86% zero-effect, ~12% negative-effect, and ~2% positive-effect.

How do our estimates of the corresponding values for the clock module compare with these values? If we select values of $\lambda = 0.41$ and $\delta = 1.42$, we can calculate the corresponding values for the probability of mutations in the N gene remaining within its wild-type oscillatory phenotype ('zero effect': #7), leaving for phenotypes with decreased DNA binding ('negative effect': #6, #8), and leaving for phenotypes with increased DNA binding ('positive effect': #3, #11). The resulting values for the distribution are then ~ 86% zero-effect, ~12% negative-effect, and ~2% positive-effect, which matches the experimental results.

The mechanism-specific *lac* system and the mechanism-nonspecific *Chlamydomonas* system are very different, but if one considers the relative frequency of the 'zero effect' mutants as an estimate of the selective pressure on the system in the selecting condition, then we can speculate that the selection for the LAC repressor is less than that for the growth rate of the entire organism. Moreover, this suggests that an experimental estimation of the distribution can be used to fit values for the $\lambda$ and $\delta$ parameters that can then be used in the population model to back calculate a numerical value for the strength of the selection (see **Figure 7**).

##### ***S12. Temporal Response at Higher Mutation Rates***

At high mutation rates the equilibrium distributions are nearly identical regardless of whether there are neutral or protein burden fitness effects in the non-selecting condition (**Figure 3A vs. 3B**). This near identity is also observed during the dynamic transition from the equilibrium distributions under non-selecting to that under selecting conditions: neutral fitness effects (**Figures S8A**) and protein burden fitness effects (**Figure S8B**). In both cases, the frequency of

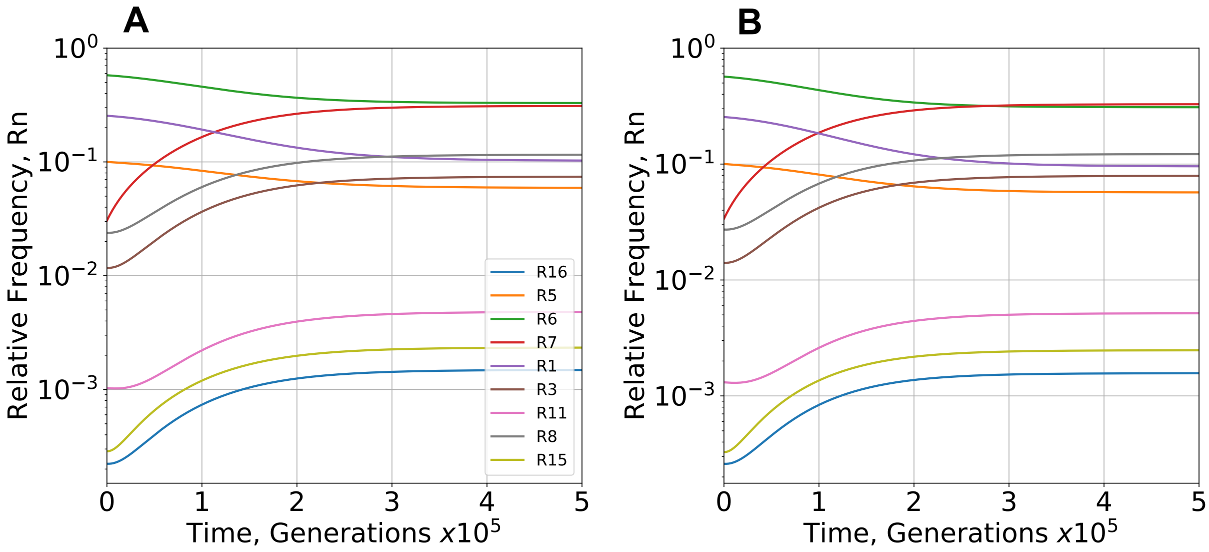


**Figure S8. Temporal response in relative frequency of phenotypes following imposition of the selecting condition.** Imposition occurs by a change from a non-selecting ($\mu_7^*$ = 1.0) to a selecting ($\mu_7^*$ = 1.00006) environment. **(A)** Neutral and **(B)** protein burden fitness effects were considered with a general mutation rate $m = 10^{-4}$. See also **Figure 5A&C**.

the selected wild-type (oscillatory phenotype #7) increases, the three highest entropy phenotypes (#1, #5 and #6) exhibit a decrease, while the other five phenotypes actually exhibit an increase along with the selected phenotype because of differences in phenotype-specific mutation rates. This result resembles genetic hitch-hiking (Smith & Haigh, 1974), but the mechanism is entirely different.
